# Enhancer plasticity in endometrial tumorigenesis demarcates non-coding somatic mutations and 3D-genome alterations boosting the oncogenic driver ESR1

**DOI:** 10.1101/2023.07.25.550368

**Authors:** Sebastian Gregoricchio, Aleksandar Kojic, Marlous Hoogstraat, Karianne Schuurman, Suzan Stelloo, Tesa M. Severson, Tracy A. O’Mara, Marjolein Droog, Abhishek A. Singh, Dylan M. Glubb, Lodewyk F.A. Wessels, Michiel Vermeulen, Flora E. van Leeuwen, Wilbert Zwart

## Abstract

The incidence and mortality of Endometrial Cancer (EC) is on the rise. 85% of ECs depend on Estrogen Receptor alpha (ERα) for proliferation, but little is known about its transcriptional regulation in these tumors.

We generated epigenomics, transcriptomics and Hi-C datastreams in healthy and tumor endometrial tissues, identifying robust ERα reprogramming and profound alterations in 3D genome organization that lead to a gain of tumor-specific enhancer activity during EC development. Integration with endometrial cancer risk single-nucleotide polymorphisms, as well as WGS data from primary tumors and metastatic samples revealed a striking enrichment of risk variants and non-coding somatic mutations at tumor-enriched ERα sites. Through machine learning-based predictions and interaction proteomics analyses, we identified an enhancer mutation which alters 3D genome conformation, impairing recruitment of the transcriptional repressor EHMT2/G9a/KMT1C, thereby alleviating transcriptional repression of *ESR1* in EC.

In summary, we identified a complex genomic-epigenomic interplay in EC development and progression, altering 3D genome organization to enhance expression of the critical driver ERα.

## Introduction

Endometrial cancer (EC) is the second most-common gynecological cancer with over 417,000 new cases and 97,000 deaths in 2020, with global incidence rates increasing every year^1^. Surprisingly, and in stark contrast to most other cancers, survival rates for endometrial cancer are decreasing^1,2^. This gradual deterioration of EC survival is likely related to the relatively understudied nature of the disease, with many molecular mechanisms driving tumor development and progression remaining largely elusive.

Endometrial tissue is under tight endocrine control, and this remains the case upon tumorigenesis. The majority of endometrial cancers (85%) are classified as low grade endometrioid tumors, expressing Estrogen Receptor alpha (ERα) and Progesterone Receptor (PR)^3–5^. PR agonists are used in the treatment of endometrial tumors as alternative to chemotherapy and hysterectomy to retain uterine function in young patients^6^. Furthermore, the estrogen competitive ERα antagonist tamoxifen is also prescribed to patients with endometrial cancer, often alternated with PR agonists (reviewed in ref ^7^). Thus, while PR acts in a tumor suppressive manner, ERα serves as driver of tumor progression which is therapeutically blocked in endometrial cancer care.

ERα is an enhancer-acting transcription factor, regulating expression of its responsive genes through long-range 3D genome interactions^8,9^. To date, most literature on promoter-enhancer communication and ERα activity has been focused on breast cancer^9^, and is far less understood in endometrial tumors. Between breast cancer and endometrial cancer, substantial differences are observed in ERα chromatin binding profiles^10–12^, highlighting the highly context-dependent nature of ERα genomic action. Paradoxically, while tamoxifen can be prescribed in the treatment of endometrial cancer, its use in the treatment of breast cancer is also reported as risk factor for endometrial tumorigenesis^13^. Previously, we showed that tamoxifen treatment in endometrial tissue reprograms the ERα chromatin interaction landscape, phenocopying profiles found in breast cancer cells, driving endometrial tumor growth^14^. These observations highlighted that the endometrial cancer epigenome is not fixed, but rather dynamically affected by oncogenic factors.

Both ERα in breast tissue^15^, and Androgen Receptor (AR) in prostate tissue^16–18^, undergo extensive reprogramming throughout the genome during tumorigenesis. In case of prostate cancer, both somatic mutations and risk single-nucleotide polymorphisms (SNPs) are enriched at tumor-gained AR sites, with only a small fraction of which causally affecting transcriptional output^17^. In endometrial cancer, such studies have thus far not been reported, and functional interplay between somatic mutations, epigenetic alterations and the impact on 3D genome organization remain fully elusive.

Here, we investigated the plasticity of ERα genomic action upon endometrial tumor development, by comparing the epigenome of primary human healthy and tumor endometrial tissues (ERα and H3K27ac ChIP-seq) and the effects on 3D genome organization (Hi-C, 4C-seq and, H3K27ac HiChIP). Secondly, we studied the crosstalk between epigenetic alterations and somatic variant events (WGS) that are associated with EC progression. We identified substantial epigenetic reprogramming upon tumorigenesis that results in the ERα re-localization at tumor-specific regions throughout the genome, which coincided with the occurrence of somatic variants in metastatic samples. In particular, we discovered an EC-specific *ESR1* enhancer which is selectively found mutated in metastatic EC. *In vitro* analyses show diminished capacity to bind lysine methyl-transferase G9a/EHMT2/KMT1C to this region when mutated, and perturbation of G9a expression enhanced ERα expression in cell line studies. Cumulatively, we show that non-coding mutations in endometrial cancer may have direct pro-tumorigenic potential, through a tight interplay with epigenetic alterations and changes in the 3D genome structure.

## Results

### ER**α** enhancer plasticity in endometrial tumorigenesis

To investigate the role of ERα signaling in endometrial tumor development, we first sought to assess how the ERα cistrome differs between endometrial tumors and healthy endometrial tissue. We collected five fresh-frozen endometrial tumors from post-menopausal patients that did not receive any estrogen receptor targeting for cancer therapy, as well as four fresh-frozen samples from post-menopausal women with pathologically normal endometrial tissue (**Fig. 1a** and **Supplementary Fig. 1a**, for clinicopathological characteristics see **Supplementary Table 1**). All samples were subjected to ERα and H3K27 acetylation (H3K27ac) ChIP-seq in order to identify active regulatory elements that are under ERα control. On average, we identified 20,815 ERα peaks (range 1,399-51,888) and 44,824 H3K27ac peaks (range 8,791-71,898) in our samples. The ChIP-seq library size was comparable among the different samples for both targets (**Supplementary Fig. 1b**). Interestingly, the number of ERα peaks was higher in tumor samples compared to healthy tissue (Mann-Whitney *P* = 0.063, **Fig. 1b**) while the global number of H3K27ac peaks for these samples was comparable between both groups (Mann-Whitney *P* = 0.29, **Fig. 1c**). To assess data quality and reproducibility, we performed Principal Component Analysis (PCA) on the peak intensities which revealed a clustering of the samples by ChIP target (first component) and tissue type (second component) (**Supplementary Fig. 1c**). Furthermore, given the relatively small sample size, we tested for over-training and performed unsupervised permutation clustering analyses that showed a clear separation of the two tissues types, except for the sample T33 showing mixed features, between normal and tumor tissues (**Supplementary Fig. 1d**), possibly due to a lower tumor cell representation in those sections that were used for ChIP-seq studies.

**Fig. 1:**
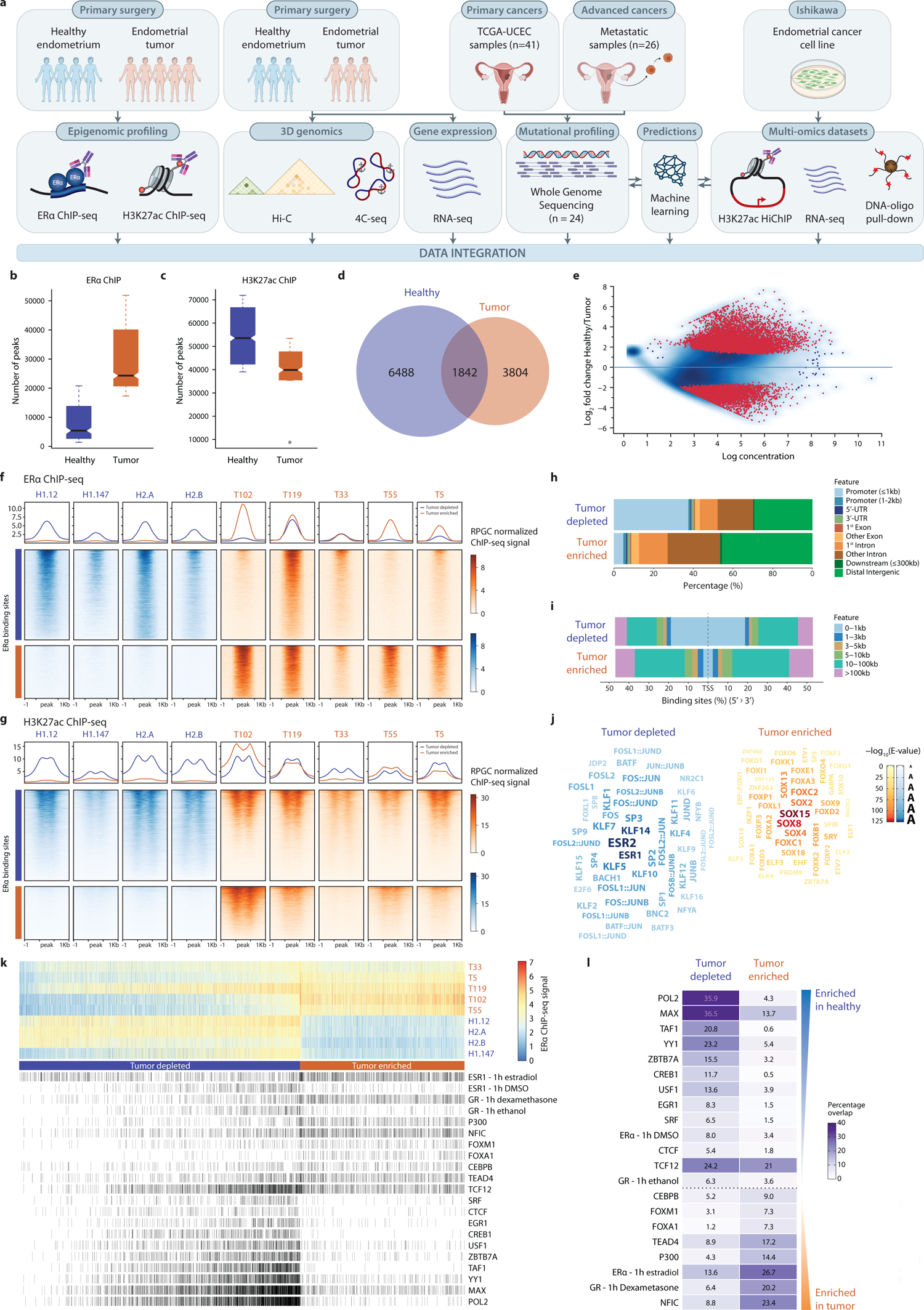
ER_α_ cistrome changes upon tumorigenesis. **a** Schematic workflow of the multi-omics approach applied in this work. **b-c** Boxplot depicting the distribution of peak number detected by ERα (**b**) or H3K27ac (**c**) ChIP-seq in healthy (blue) or tumor (orange) endometrial tissues. **d** Venn diagram of the overlap between consensus ERα binding sites detected in healthy (blue) and tumor (orange) tissues. **e** MA-plot of differential binding analyses performed on the consensus ERα ChIP-seq peaks (Healthy vs Tumor). Differential binding sites (FDR ≤ 0.05 & |log_2_(FoldChange)| ≥ 1) are highlighted in pink. **f-g** Tornado plot of the ERα (**f**) or H3K27ac (**g**) ChIP-seq signal at tumor-depleted (blue, upper blocks) or tumor-enriched (orange, lower blocks) ERα consensus peaks for all the 4 healthy (blue, left heatmaps) and 5 tumor (orange, right heatmaps) endometrial tissues. On the top of each heatmap is plotted the average density signal for each region group. **h** Stacked bar plot depicting the genomic localization frequency of tumor-depleted and tumor-enriched ERα ChIP-seq consensus peaks. **i** Stacked bar plot displaying the distance to associated gene TSS frequency distribution of tumor-depleted and tumor-enriched ERα ChIP-seq consensus peaks. **j** Wordcloud of the top-50 enriched motifs at tumor-depleted (blues) and tumor-enriched (oranges) ERα consensus peaks. Size and color intensity are proportional to the -log_10_(E-value). **k** The upper part of the heatmap shows the individual tumor-depleted (blues) and tumor-enriched (oranges) ERα consensus peaks colored by signal intensity. On the lower part, each black bar represents an overlapping peak of ChIP-seq data of several targets publicly available for the Ishikawa endometrial cancer cell line. **l** Heatmap of the percentage of tumor-depleted or tumor-enriched ERα consensus peaks overlapping with each ChIP-seq target in Ishikawa cells from (**k**). Ranking is performed by descending number of overlaps in tumor-depleted peaks.

Differential binding analysis (*DiffBind*^19^) for ERα between tumor and normal samples revealed 10,292 differentially-bound genomic locations (FDR ≤ 0.05 & |log_2_(FoldChange)| ≥ 1) (**Fig. 1d-e** and **Supplementary Fig. 1f**), of which 6,488 ERα sites were lost and 3,804 ERα sites gained in tumor samples, hereafter referred to as “tumor-depleted” (blue) and “tumor-enriched” (orange) ERα binding sites, respectively. Tornado plots depicting normalized ERα ChIP-seq signal for all 10,292 differential-enriched ERα sites (**Fig. 1f**), and label-swap permutation differential binding analyses (**Supplementary Fig. 1e**), confirmed the selective enrichment of signals between samples, with high reproducibility. As ERα is a transcription factor that acts predominantly by occupying enhancers^8,9^, we next analyzed the differential ERα sites for presence of H3K27ac, inferring regulatory element activity. Interestingly, persistent activity in both tissue states was observed at tumor-depleted Erα sites, while tumor-enriched ERα sites showed a marked increase of activity upon tumorigenesis (**Fig. 1g**). Collectively, these data suggest a selective gain of regulatory element potential upon tumorigenesis, marked by dynamically altered ERα sites.

### Genomic distributions and transcription complex composition at differential ER**α** sites

To further investigate the nature of the ERα regions differentially bound upon tumorigenesis, we first inspected their genome-wide distribution with respect to different genomic elements. In agreement with previous reports from us (tumors^10,14^) and others (cell lines^11^), tumor-enriched ERα sites were mostly found at distal intergenic and intronic regions (**Fig. 1h**) and at larger distance from the associated Transcription start site (TSS) (**Fig. 1i**). However, tumor-depleted ERα sites, selectively occupied by ERα in healthy endometrial tissue, display a strong enrichment for promoter regions of genes involved in cell cycle regulation and DNA damage processes (**Supplementary Fig. 1g**). To determine whether loss of ERα binding at these promoters leads to transcriptional deregulation of the associated genes in tumor samples, we performed RNA sequencing (RNA-seq) in 3 healthy and 3 tumor endometrial tissues from post-menopausal patients that did not receive any prior estrogen receptor targeting for cancer therapy (**Supplementary Table 1**). Principle component analyses display a clear separation of normal and tumor tissues with a higher variability for tumor samples than healthy ones (**Supplementary Fig. 2a**). Differential expression analyses identified 1945 genes to be down-regulated and 2392 to be up-regulated in tumors, compared to healthy tissue (**Supplementary Fig. 2b**). Next, we analyzed the expression of the tumor-depleted ERα promoter-bound genes included in the previously identified over-represented biological processes (**Supplementary Fig. 1g** and **Supplementary Fig. 2c**). This investigation confirmed our former results highlighting a significant deregulation of cell division and DNA-repair related genes (**Supplementary Fig. 2c**). Furthermore, results from Gene Set Enrichment Analyses (GSEA) for the “hallmarks” (H) dataset also corroborated with these findings in a more global manner (**Supplementary Fig. 2d**). Of note, we observed that differentially expressed genes associated with tumor-enriched ERα binding sites were mostly up-regulated in tumor samples compared to genes associated with tumor-depleted ERα binding sites (Fisher’s exact *P* < 2.2×10^-16^, OR 6.8, 95% CI 4.35-10.84; **Supplementary Fig. 2e-f**).

Consensus motifs for ERα/β (ESR1/2) and AP-1 (JUN, FOS) binding sites were specifically enriched at tumor-depleted ERα regions, whereas tumor-enriched regions demonstrated an overabundance of SOX and FOX transcription factor family member motifs (**Fig. 1j**). Interestingly, tumor-enriched ERα binding sites, despite the consensus ERα/ESR1 motif being poorly detected, do harbor this motif in around 85% of the cases in a measure comparable to the tumor-depleted sites (**Supplementary Fig. 1h**), albeit highly degenerated as compared to tumor-depleted or common ones (**Supplementary Fig. 1i**). Differential motif enrichment analyses led us to hypothesize that the ERα transcription complex may differ between tumor-enriched and tumor-depleted sites. To test this hypothesis, we next integrated our tumor-enriched and tumor-depleted sites with publicly available ChIP-seq data from the Ishikawa endometrial cancer cell line (**Fig. 1k** and **Supplementary Table 2**), revealing striking differences between transcription and epigenetic factors on chromatin occupancy for the two ERα binding site categories. Interestingly, ERα chromatin binding in hormone-deprived cells showed a significant overlap with the tumor-depleted ERα peaks (adjusted Fisher’s exact *P* = 3.15×10^-21^), while ERα peaks from estradiol-stimulated cells showed significant overlap with the tumor-enriched ERα peaks (adjusted Fisher’s exact *P* = 4.60×10^-59^, **Fig. 1l**). In addition, both FOXM1 and FOXA1 binding sites in Ishikawa showed significant overlaps with our tumor enriched regions (adjusted Fisher’s exact *P* = 3.37×10^-21^ and *P* = 6.25×10^-57^, respectively, **Fig. 1l**). These findings were successfully confirmed by *GIGGLE*^20^ analyses (testing for overlap in a large repository of thousands of publicly available ChIP-seq datasets) at differential ERα binding sites (**Supplementary Fig. 1j**) implicating a strongly divergent transcription complex composition between tumor-enriched and tumor-depleted ERα sites. Altogether, these data suggest prominent enhancer plasticity in endometrial tumorigenesis, with ERα relocating to alternatively engaged and activated non-canonical enhancers in endometrial cancer.

### 3D high-order chromatin structures are altered in tumors

Enhancer regions can modulate gene transcription through interactions with promoters in 3D genomic space^21^. With the observed plasticity of ERα chromatin profiles in tumorigenesis, we hypothesized that these epigenetic alterations were accompanied by reorganization of the 3D chromatin structure. To test this hypothesis, we performed high-throughput chromosome conformation capture (Hi-C) analyses on healthy endometrial tissue (n=3) and primary tumors (n=3) derived from post-menopausal women that did not receive any previous systemic therapy (**Fig. 2a**, for clinicopathological features see **Supplementary Table 1**). Hi-C is particularly suitable to study translocations in cancer^22^, which were not observed in any of the tumor samples we analyzed (**Supplementary Fig. 3a**). We did however observe a distinct clustering of the healthy tissues from the tumor specimens, based on 3D genome organization (**Supplementary Fig. 3b**) and independent of Hi-C contact quality bias, intrinsic to the tissue type (**Supplementary Fig. 3c**). In addition, tumor samples displayed a higher Hi-C contact heterogeneity in comparison to the more correlated healthy tissues (**Supplementary Fig. 3c**). Next, we quantified the Relative Hi-C Contact Probability (RCP) as function of the contact distances. Interestingly, tumors displayed an increased probability of shorter-range interactions (<2Mb) relative to healthy tissue, which is concomitant with a loss of longer-range chromatin contacts (**Fig. 2b-d** and **Supplementary Fig. 3d-e**), highlighting a major 3D genome reorganization in tumor samples independently of the inter-individual variability. Furthermore, analyzing the distribution of the loop distances (**Fig. 2e** and **Supplementary Fig. 4**), we found that loops whose anchors include tumor-enriched ERα binding sites tend to be shorter in tumor tissues than in healthy tissues. On the other hand, tumor-depleted and shared ERα binding sites do not show differences in loop length between two tissues stages. We therefore investigate the distribution of the Hi-C contacts as function of the distance from different categories of ERα binding sites, by performing paired-end spatial chromatin analysis (PE-SCAn^23^) between healthy and tumor tissues (**Fig. 2f**). We observed that shared and tumor-depleted ERα binding sites lose chromatin interaction in their surrounding regions in tumor tissues, whereas tumor-enriched binding sites show an increase in relatively shorter-range interactions (**Fig. 2f**). We next wondered whether the loop shortening, and increased chromatin contacts, at tumor-enriched binding sites in tumor samples was associated with changes in chromatin compartmentalization. Analysis of compartment polarization (**Fig. 2b,g-i** and **Supplementary Fig. 3f-g**) revealed decreased compartment strength in tumor samples compared to healthy tissues. In particular, we observed that A compartments (euchromatin) were more robust in healthy samples, while B compartments (heterochromatin) were slightly stronger in tumor tissues (**Fig. 2h** and **Supplementary Fig. 3g**). Overall, around one third of compartments (A: 565/1431 (39.5%); B: 440/1306 (33.6%)) underwent class switching in tumor samples (**Fig. 2i**). To investigate whether this compartment class switch was occurring specifically in compartments where ERα was bound in a tissue type-specific manner, we overlapped the compartments with the differential ERα occupied regions upon tumorigenesis (**Fig. 2j**). We found that compartments switching from A to B class were enriched for tumor-depleted ERα binding sites, relative to tumor-enriched ones (A-to-B depleted/enriched ratio 2.65 (135/51)). In contrast, compartments switching from B-to-A were clearly enriched for tumor-gained ERα binding sites (B-to-A depleted/enriched ratio 0.33 (35/105)). These findings were further confirmed studying the behavior of smaller genomic regions (compartment bins) relative to differentially-occupied ERα binding sites, excluding regions that harbor multiple ERα binding site categories (Fisher’s exact test: B-to-A depleted-vs-enriched: *P* = 1.317×10^-56^, OR 0.20; A-to-B depleted-vs-enriched: *P* = 6.013×10^-11^, OR 1.87) (**Fig. 2k** and **Supplementary Fig. 3h**).

**Fig. 2:**
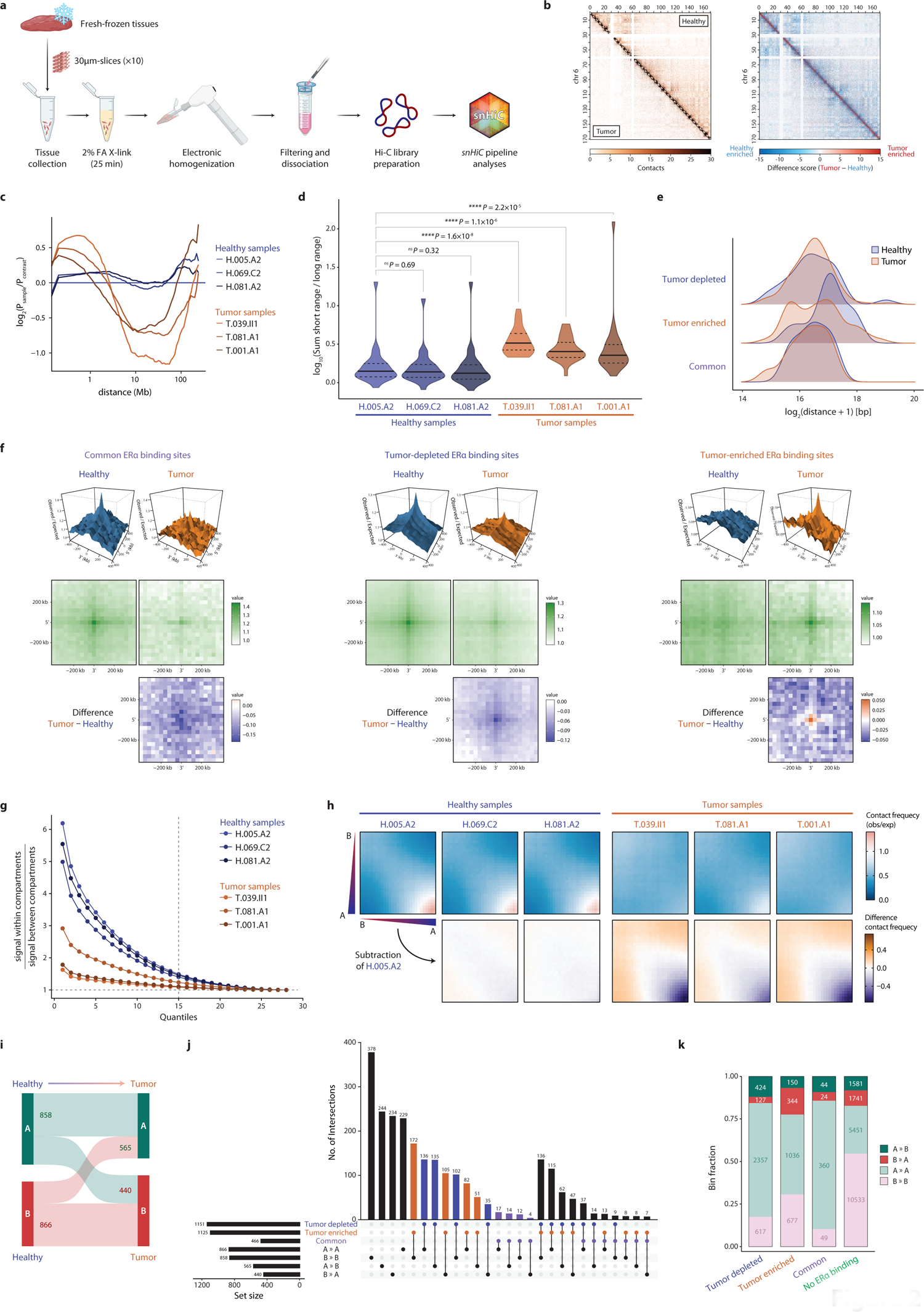
3D genome landscape is remodeled during tumorigenesis. **a** Schematic of the Hi-C library preparation starting from fresh frozen tissues. Ten 30µm-thick slices of flash-frozen tissue are cross-linked 25min in 2% formaldehyde. Then, electronically homogenized tissues are filtered using a 75µm cell strainer and subjected to Hi-C library preparation. Hi-C libraries are then sequenced and resulting reads are analyzed by the *snHiC*^68^ pipeline. **b** On the left, 40kb-resolution matrices of the average Hi-C score in Tumor (n = 3) and Healthy endometrial tissues (n = 3) at chromosome 6. On the right, differences of the scores shown on the left side (Tumor – healthy). **c** Relative Hi-C Contact Probability (RCP) as function of the distance for each individual sample (40kb resolution). **d** Violin plot of the distribution of the short-range (<2Mb) over long-rage (>2Mb) Hi-C contacts ratio in each individual sample (40kb resolution). For each comparison the Wilcoxon’s test *P*-value is indicated. **e** Distribution of the Hi-C loop distance (10kb resolution) for loops overlapping with tumor-depleted, tumor-enriched or shared ERα consensus peaks in combined healthy (blue) or tumor (orange) endometrial tissues. **f** PE-SCAn results in healthy and tumor tissues at common, tumor-depleted and tumor-enriched ERα binding sites. For each panel, top and middle rows depict the 3D and 2D, respectively, representation of the Hi-C contact frequency distribution in healthy (left) and tumor (right) tissues; lower row shows the difference of Hi-C contact frequencies between Tumor (orange) and Healthy (blue) tissue scores. **g** Compartment polarization ratio (100kb resolution), defined as (AA + BB) / (AB + BA), for each individual sample. **h** *Top*: saddle plot of A/B compartments interactions as computed in (**g**) for each individual sample. *Bottom*: difference of saddle-score with the reference H.005.A2 (healthy tissue), where orange indicates a higher score in tumor samples while purple indicates a higher score in healthy tissues. **i** Sankey plot depicting the proportion of compartment state transition (100kb resolution) of healthy compartments (left) upon tumorigenesis (right). **j** Upset plot showing the overlaps between different compartment transition states (100kb resolution) in healthy compared to tumor as in (**g**) (A-to-A, B-to-B, A-to-B, B-to-A) and different categories of ERα consensus peaks. **J** Stacked bar plot of the proportion of compartment transition status as in (**g**) for individual genomic bins overlapping with only one of the different categories of ERα consensus peaks.

Taken together, our findings highlight a widespread reorganization of the 3D genome upon tumorigenesis that leads to a loss of chromatin compartment polarization. This was accompanied by an extensive class-switching, with tumor-enriched ERα binding sites being enriched in B-to-A compartments. Moreover, we identified a gain of relatively shorter-range chromatin interactions upon tumor development, at the expense of longer-range contacts.

### Somatic mutations are selectively found at a higher frequency at tumor-enriched ER**α** binding sites

Recently, pan-cancer whole-genome analysis highlighted the presence of driver mutations in non-coding regulatory elements^24^. To determine whether that might also be the case in endometrial cancer, we re-analyzed Whole-Genome Sequencing (WGS) data of 41 primary endometrial cancer samples from the TCGA Uterine Corpus Endometrial Carcinoma (UCEC) cohort^25^ (see Methods for details on the cases selection). Most of the mutations were located in intronic and intergenic non-coding regions (**Fig. 3a**). Mutational signatures (**Fig. 3b**) and variant counts (**Fig. 3c**) well correlated with the Micro-Satellite Stability (MSS) status of the source samples. These analyses show that most of the Micro-Satellite Instable (MSI) samples are characterized by UV light exposure (SBS7c) and defective DNA mismatch repair (SBS15/44) signatures (**Fig. 3b**), accompanied by a higher number of mutations per sample (**Fig. 3c**). On the other hand, Micro-Satellite Stable (MSS) samples, as expected, display a lower number of somatic mutation counts (**Fig. 3c**) and prevalently mutational signatures associated to defective DNA (polymerase ε exonuclease domain mutations, SBS10b) (**Fig. 3b**).

**Fig. 3:**
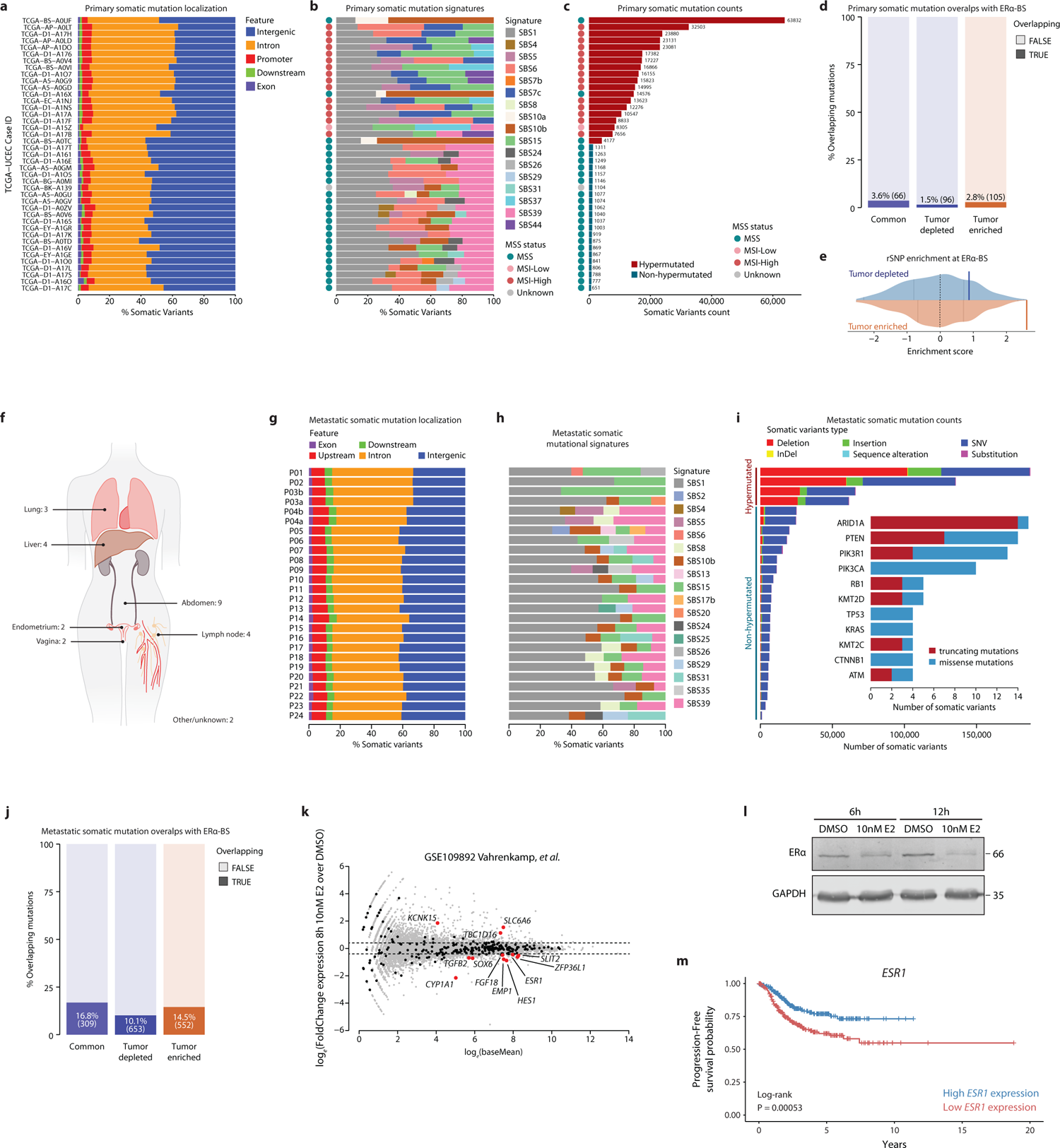
Tumor-enriched ER_α_ binding sites represent non-coding regions target of somatic mutation in metastatic endometrial cancer tumor. **a-c** Genomic localization (**a**), mutational signature (**b**) and counts (**c**) distribution of the somatic mutations detected by WGS in the primary endometrial cancer samples form a selection of samples from the TCGA-UCEC cohort (n=41). **d** Stacked bar blot depicting the fraction of ERα peaks overlapping with somatic mutations identified in the primary cohort. **e** Variant Set Enrichment (VSE) analysis depicting the enrichment of differentially bound ERα sites over endometrial cancer risk loci identified by genome-wide association study (GWAS, *P* < 1×10^-5^)^26^. Density plot represents distribution for the Z-score from matched controls defining the null distribution. Blue (tumor-depleted) or orange (tumor-enriched) vertical lines represent observed enrichments, with tumor-enriched enrichment being statistically significant (*P_adj_* = 0.0060). Gray vertical lines define 0, 25, 75, 100 percentiles of the distribution, while dotted black line indicates the median. **f** Scheme of the number metastases and metastatic site, for all metastatic samples used for the WGS analyses. **g-h** Genomic localization (**g**) and mutational signature (**h**) distribution of the somatic mutations detected by WGS in the metastatic samples described in (**f**). **i** In the outer plot, it is shown the number of different types of somatic variants detected in each metastatic sample. In the inner plot, stacked bar plot of the most-frequently protein sequence mutated genes and relative number and type of somatic variants identified in metastatic samples. **j** Stacked bar blot depicting the fraction of ERα peaks overlapping with somatic mutations identified in the metastatic cohort. **k** MA-plot of differential expression analyses in Ishikawa cell lines upon 8h of 10nM β-estradiol (E2) stimulation. Black dots indicate genes which promoter has been linked, by H3K27ac Hi-C analyses, to a tumor-depleted or tumor-enriched ERα-bound regulatory element bearing somatic mutations in metastatic samples (n=311). Differentially expressed genes (|Fold Change expression| ≥ 1.5 & *P*_adj_ < 0.05) upon estradiol stimulation have been labeled and highlighted by a red dot (n=12). **l** Ishikawa endometrial cancer cells were treated or not for 6 or 12 hours with 10nM estradiol. Whole-cell extracts were analyzed by immunoblotting with antibodies against ERα and GAPDH (loading control). **m** Progression-Free Kaplan-Meier curve of endometrial cancer patients (TCGA data) divided into two groups using the median of *ESR1* expression (FPKM) as cut-off.

To investigate whether somatic mutations in primary tumors are associated with selective ERα activity, we compared the number of somatic mutations that are overlapping with the different categories of ERα binding sites. We found that tumor-enriched binding sites have a higher representation of somatic mutations over the tumor-depleted ones (Fisher’s exact *P* = 1.143×10^-5^, OR 1.89, 95% CI 1.41-2.53; **Fig. 3d**). Notably, the somatic variants overlapping with the ERα binding sites are carried almost exclusively by hyper-mutated samples (**Supplementary Fig. 6a**). Interestingly, analyzing the enrichment of Genome-Wide Association Study (GWAS) loci associated with endometrial cancer risk at ERα-binding sites^26^, we observed that tumor-enriched binding sites are significantly enriched (*P_adj_* = 0.006) for endometrial cancer risk single-nucleotide polymorphisms (rSNP) in comparison to tumor-depleted regions (*P_adj_* = 0.995) (**Fig. 3e**). These data imply that both germline as well as somatic variants are not equally distributed over the genome, but rather show a selective occurrence at tumor-gained ERα sites.

To investigate whether somatic variant enrichment for particular ERα sites also persisted after endometrial cancer progression, we next analyzed WGS data of 26 fresh frozen biopsy samples, derived from 24 cases of advanced endometrial cancer. Biopsies were collected from tumors that metastasized to lung, liver, and other abdominal regions (**Fig. 3f** and **Supplementary Table 3**). In these tumors, the vast majority (>96%) of variants were located at non-coding regions, with 85% of the identified variants found to populate intronic and intergenic regions. Considerably, less variants (mean 9%, range 8.1-10.3) were located proximally upstream of a coding gene, potentially representing promoter regions (**Fig. 3g**). To assess WGS data quality and explore additional genomic features of our samples, we investigated mutational signatures, counts, and frequently-mutated genes (**Fig. 3h-i**). Four samples were hypermutated and displayed 20-67% of variants associated with defective DNA mismatch repair (SBS15). Six samples from tumors pre-treated with carboplatin showed traces of signatures 31 and 35, both associated with platinum treatment^27^. Cumulatively, these results confirm previously-described enrichment of tumor-intrinsic^28,29^ and treatment-induced^27^ mutational features of endometrial cancer. The most-frequently mutated genes with protein sequence altering mutations included *ARID1A* and various members of the PIK3CA pathway, such as *PTEN*, *PIK3CA* and *PIK3R1*, recapitulating previous observations for endometrial tumor^28^.

As expected, analogously to the primary tumor analyses (**Fig. 3a**), somatic mutations in metastatic samples were also predominantly occurring in non-coding regions. Next, we integrated these somatic mutation data with tissues-type enriched ERα ChIP-seq data (**Fig. 3j**), and again found differentially bound ERα regions were significantly enriched for somatic mutations (11.9%, 1,221/10,292), relative to the total number of ERα-bound sites from our entire cohort, not found to differ between normal tissue and tumors (8%, 5,881/73,312; Fisher’s exact *P* = 5.64×10^-36^, OR 1.54, 95% CI 1.44-1.65). All samples contributed to this enrichment, with a median somatic variant count of 18 (range 1-358) in differentially ERα-bound sites. Moreover, comparing tumor-enriched with tumor-depleted regions, we observed a significant enrichment of somatic variants in the tumor-enriched ERα sites compared to tumor-depleted ones (14.5% vs 10.1%, Fisher’s exact *P* = 2.98×10^-11^, OR 1.51, 95% CI 1.34-1.71, **Fig. 3j**). In contrast to primary tumors, these observations are independent of the mutational frequency status (**Supplementary Fig. 6b**). In most instances (n=1,002), regions overlapped with only one somatic variant, while some ERα-bound regions were observed to harbor 2 (183 regions), 3 (34 regions) or 4 (2 regions) somatic mutations.

To investigate genes whose expression may be affected by enhancer mutations, we intersected tumor-depleted and tumor-enriched ERα binding sites (FDR ≤ 0.01) bearing mutations with H3K27ac HiChIP chromatin looping data in Ishikawa cells (**Supplementary Table 4**), aimed to couple putative enhancers with the transcription start sites (TSS) they control, yielding a total of 331 genes. Out of these, 13 genes were differentially expressed in Ishikawa endometrial cancer cells upon 8h of estradiol-stimulation^30^ (**Fig. 3k**). Interestingly, *ESR1* (encoding ERα) was represented as well, being modestly downregulated at both RNA and protein level (**Fig. 3k-l** and **Supplementary Fig. 7c**), and harboring 3 metastatic somatic mutations in two *cis* regulatory elements upstream of the *ESR1* locus, hereafter referred as ‘*Enhancer 1*’ (P17 and P22; non-hypermutated) and ‘*Enhancer 2*’ (P03b; hypermutated) (**Fig. 4a** and **Supplementary Fig. 7b**). Of note, high *ESR1* expression is associated with favorable outcome in primary endometrial cancer patients (TCGA^25^) (**Fig. 3m**). Of note, these two enhancers are specific to the endometrium cancer samples, relative to healthy endometrial tissues and breast cancer specimens, based on ATAC-seq (TCGA^31^) and ERα ChIP-seq signal (our previous work^32^) (**Supplementary Fig. 7b-c**) at the *ESR1* locus.

**Fig. 4:**
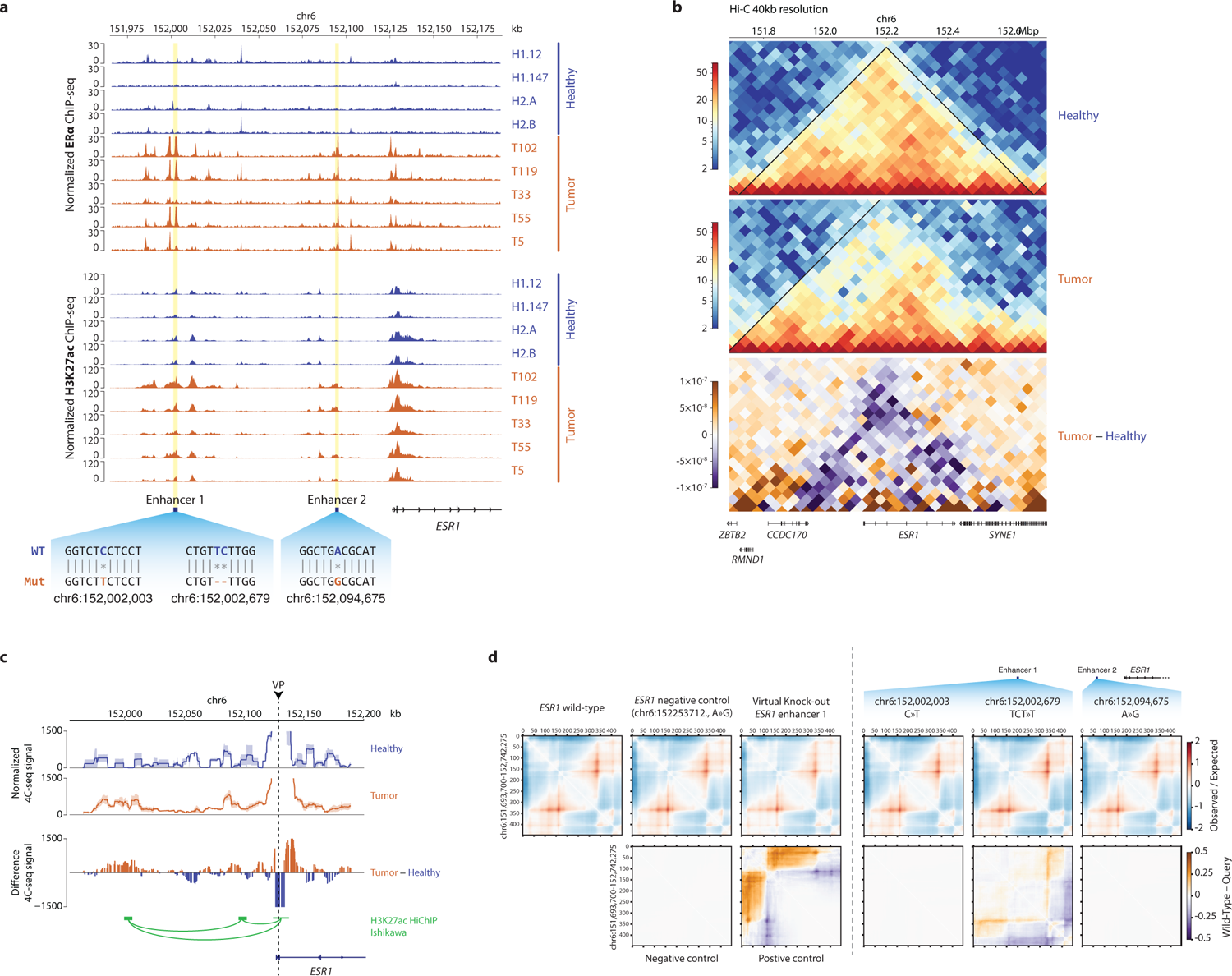
Short-range chromatin contacts at the *ESR1* locus are stronger in tumors. **a** ChIP-seq genomic tracks for ERα (upper panel) and H3K27ac (lower panel) in healthy (blue) and tumor (orange) endometrial primary tissues at the *ESR1* locus. *ESR1 Enhancer 1* and *Enhancer 2* are indicated, as well as the mutations found by WGS analyses in metastatic samples. **b** Representation of the averaged 40kb resolution Hi-C matrix at the *ESR1* locus for the 3 healthy (top) and 3 tumor (middle) tissues or the score difference (bottom) tumor – healthy, where orange indicates higher scores in the tumors while purple higher scores in healthy. The matrices scores are dived by the sum of the matrix. Black lines indicate the Topologically Associated Domains (TADs) identified. **c** 4C-seq genomic tracks at the *ESR1* locus using the *ESR1* TSS as View-Point (VP) for 2 healthy (top, blue) and 3 tumor (middle, orange) endometrial tissues, and the average difference of score tumor – healthy (bottom). With green arcs are depicted loops detected by H3K27ac HiChIP in Ishikawa endometrial cancer cells. The ribbon around the 4C-seq signal lines indicates the Standard Error Mean (SEM) among biological replicates. **d** In the top row, Observed/Expected matrices of sequence-based machine learning prediction in a ±500kb window surrounding the *ESR1* locus. In order from left to right can be found: wild-type sequence, point mutation in a genomic desert (negative control), deletion of the full *Enhancer 1* sequence (positive control), introduction of SNVs found by WGS analyses of metastatic endometrial cancer samples. On the bottom row, is showed the difference of Observed/Expected score over the wild-type sequence, where orange indicates higher scores in the altered sequence and purple a higher score in the wild-type one.

### *ESR1* enhancer mutation alters 3D genome contacts and decreases EHMT2/G9a enhancer binding to boost ERα expression

Since *ESR1* expression is strongly associated with favorable outcome in endometrial cancer (**Fig. 3h**), alterations in its transcriptional regulation may have direct implications on tumor development and progression. Since both *Enhancer 1* and *Enhancer 2* – found mutated in metastatic, but not primary endometrial cancer – showed induction of ERα binding and H3K27ac signals in tumors (**Fig. 4a**), we investigated whether these loci may undergo alternative regulation in different stages of tumor progression leading to *ESR1* expression deregulation. Of note, in both our (**Supplementary Fig. 5a**) and TNM-plot^33^ (**Supplementary Fig. 5b**) RNA-seq expression data, *ESR1* expression in primary tumors was comparable to the levels found in normal tissue, suggesting that altered regulatory potential alone at these enhancers was not sufficient to affect *ESR1* expression.

Hi-C analyses of endometrial tumors (**Fig. 4b**) illustrated increased short-range and decreased long-range chromatin interactions at this locus, when compared to healthy endometrial tissue. These data were confirmed with 4C-seq analyses using the *ESR1* promoter as view-point (**Fig. 4c**), showing tumor-specific gained interactions of newly ERα-engaged enhancers with the *ESR1* promoter (**Supplementary Fig. 3i**).

To identify which *ESR1* enhancer mutation had an higher probability to impact *ESR1* gene regulation and potentially drive tumor progression, we employed the *Akita*^34^ machine learning tool to predict the effect of the somatic mutations on the 3D genome organization (**Fig. 4d**). Our analyses predicted the mutation occurring at position chr6:152,002,679 (TTC-to-T, P22) of *Enhancer 1* to affect chromatin architecture surrounding the *ESR1* locus. Based on these results, this specific mutation was selected for further analyses.

To determine whether enhancer mutations can alter the composition of the transcription complex recruitment, we performed DNA affinity purification experiments coupled to quantitative mass spectrometry. To this end, we incubated nuclear extracts from Ishikawa endometrial cancer cells with immobilized DNA-oligonucleotides encompassing the center of the ERα-bound site at the *Enhancer 1* – containing either the reference or mutant sequence – with a ±25bp window around it (**Fig. 5a-b**). Quantitative proteomics identified 13 gained and 11 lost proteins upon mutation of the regulatory element, respectively (**Fig. 5b**). To identify differentially bound proteins that may directly regulate *ESR1* expression, we used publicly available RNA-seq (TCGA^25^) data of 589 endometrial cancer tissues to correlate the expression of *ESR1* with the expression of genes corresponding to the 24 differentially-bound proteins (**Fig. 5c**). Then, we selected the genes that most-strongly positively (*POLK*, *ZBTB21*, *XPA*, *SIN3A*) or negatively (*GTF2IRD1*, *EHMT2*, *ZNF768*) correlated with *ESR1* expression (**Supplementary Fig. 8a**) For these genes, expression was analyzed in relation to the progression-free survival probability of endometrial cancer patients (TCGA^25^) (**Fig. 5d** and **Supplementary Fig. 8b**). Interestingly, our analyses identified only one of the 7 genes analyzed – the lysine methyl-transferase EHMT2 (also known as G9a or KMT1C) – to significantly associate with a poor outcome of endometrial cancer patients (**Fig. 5d** and **Supplementary Fig. 8b**).

**Fig. 5:**
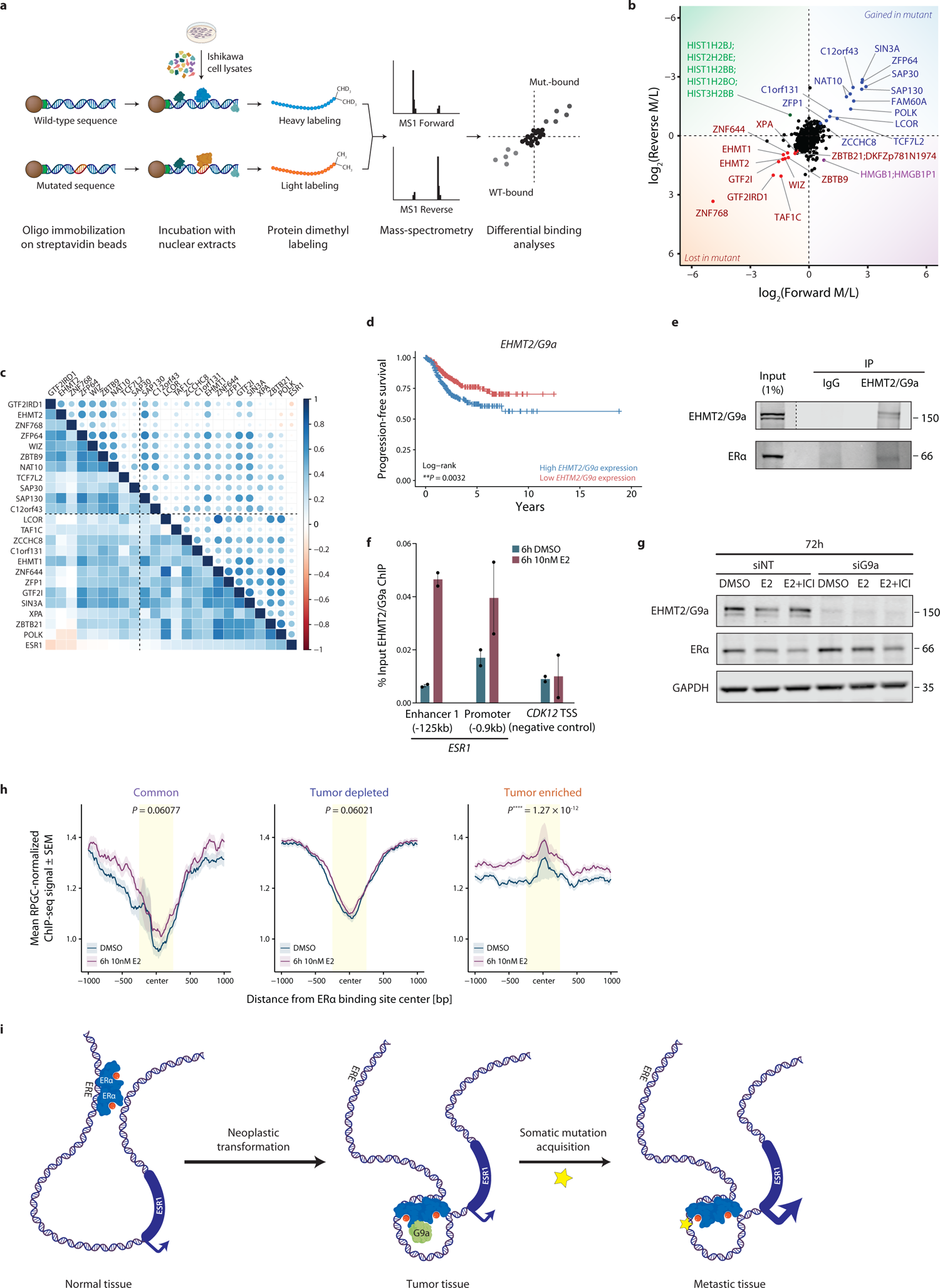
EHMT2/G9a is a negative regulator of *ESR1* expression in endometrial cancer. **a** Schematic workflow used to perform DNA-oligo protein pull-downs. Biotin-conjugated wild-type or mutated DNA-oligos are immobilized on streptavidin magnetic beads and mixed with Ishikawa nuclear lysates. Captured proteins are then dimethyl labeled and analyzed by mass spectrometry. **b** A DNA oligo with the sequence of the ±25bp surrounding the ERα binding site in the *ESR1 Enhancer 1*, in wild-type or chr6:152,002,679–TCT-to-T form, was used to perform DNA-oligo protein pull downs in Ishikawa cells as described in (**a**). The scatter plot shows the log_2_ ratios of all identified and quantified proteins in both experiments plotted against each other. Proteins significantly enriched at the wild type sequence are highlighted in red, and proteins significantly enriched at the mutant sequence are highlighted in blue. **c** Gene expression correlation heatmap for all corresponding 24 differentially bound proteins identified in (**b**) in 589 endometrial cancer patients (TCGA). Dashed lines indicate the separation between positive and negative correlation scores. Genes are ranked by correlation with *ESR1* gene expression. **d** Progression-Free Kaplan-Meier curve of endometrial cancer patients (TCGA data) divided into two groups using the median of *EHMT2/G9a* expression (FPKM) as cut-off. **e** EHMT2/G9a and IgG (negative control) ChIP followed by Western blot in Ishikawa endometrial cancer cells. Antibodies against EHMT2/G9a and ERα have been used. For EHMT2/G9a, two different exposure images were used for input and IP, as indicated by the vertical dashed bar. **f** EHMT2/G9a ChIP in Ishikawa cells stimulated for 6h with 10nM β-estradiol. Barplot shows percentage of enrichment over the input (% Input) at the *ESR1 Enhancer 1*, *ESR1* promoter and *CDK12* promoter (negative control) analyzed by quantitative PCR (qPCR). Mean of 2 independent experiments is shown. **g** Ishikawa endometrial cancer cells were stimulated for 72h with 10nM β-estradiol in combination or not with 100nM ICI-182,780 (Fulvestrant, negative control). In all conditions, cells where incubated either with a Non-Targeting (NT) siRNA or with an siRNA against *EHMT2/G9a*. Then, whole-cell extracts were analyzed by immunoblotting with antibodies against ERα, EHMT2/G9a and GAPDH (loading control). **h** Differential ERα peak centered EHMT2/G9a average ChIP-seq density profiles in Ishikawa cells upon 6h treatment with 10nM β-estradiol (purple) or DMSO (blue, control). Ribbon indicates SEM. Paired Wilcoxon test was performed on the average score of the highlighted area for each individual region analyzed; *P* values are indicated. **i** Schematic representation of the *ESR1* gene regulation working model. Upon tumorigenesis, ERα is re-located to an endometrial cancer specific *ESR1* enhancer (Fig. 1f**-g**). ERα interacts with EHMT2/G9a (Fig. 5e) and binds both enhancer and promoter regions of ESR1, in a hormone-dependent fashion (Fig. 5f). In a subset of metastatic endometrial cancers, a somatic mutation is acquired at the tumor-specific *ESR1* enhancer (Fig. 4a). *In vitro* analyses probing this mutation revealed a loss of EHMT2/G9a DNA binding capacity (Fig. 5b), and EHMT2/G9a knockdown in endometrial cancer cell lines leads to an increase of *ESR1* expression levels (Fig. 5g).

EHMT2/G9a interacts with ERα (**Fig. 5e** and **Supplementary Fig. 8c**), based on co-immunoprecipitation experiments. In line with this, ChIP-qPCR revealed that EHMT2/G9a was recruited to both *Enhancer 1* and promoter of the *ESR1* gene in Ishikawa cells, following estradiol treatment (**Fig. 5f**). As negative control, these experiments were reperformed in the ERα-negative endometrial cancer cell line AN3CA (**Supplementary Fig. 9a-b**). In line with the absence of ERα expression, estradiol treatment did not alter tumor cell growth (**Supplementary Fig. 9c-d**). In absence of ERα, there is little recruitment of EHMT2/G9a to the *ESR1 Enhancer1* and promoter when compared to the negative control region (**Supplementary Fig. 9e**). Furthermore, siRNA-mediated knock-down of EHMT2/G9a in Ishikawa cells, resulted in an increase of ERα protein expression (**Fig. 5g** and **Supplementary Fig. 8d**), confirming the reciprocal relationship between these proteins. Interestingly, we observed a slight decrease in EHTM2/G9a protein levels upon estradiol stimulation, an effect that was abolished when ERα was targeted for degradation^35^ by ICI 182,780 (Fulvestrant) treatment (**Fig. 5g** and **Supplementary Fig. 8d**). As negative control, comparable experiments performed in ERα-negative AN3CA cells, which did not show any gain of ERα protein levels upon EHMT2/G9a knock-down (**Supplementary Fig. 9f-g**).

Finally, we wondered whether the ERα-dependent chromatin recruitment of EHMT2/G9a is a specific feature of *ESR1* locus, or rather represents a more general feature of EHMT2/G9a action. To address this question, we studied the EHMT2/G9a ChIP-seq signal around the different categories of ERα binding sites in the presence or absence of estradiol stimulation in Ishikawa cells (**Fig. 5h**). These analyses revealed an absence of EHMT2/G9a signal at tumor-depleted and common regions, with no differences in its recruitment upon estradiol stimulation. In contrast, EHMT2/G9a co-localizes with ERα at tumor-enriched binding sites with a significantly increased signal upon estradiol stimulation (**Fig. 5h**, *right*), suggesting a more general behavior of EHMT2/G9a for tumor-gained ERα sites.

Altogether, these data show that EHMT2/G9a DNA binding is impaired upon mutation of the *ESR1 Enhancer 1* locus, thereby enhancing expression of tumor driver ERα in metastatic endometrial cancer and that its activity is required to repress *ESR1* expression in endometrial cancer in a ERα-dependent manner.

## Discussion

High expression levels of *ESR1* (encoding ERα) in endometrial cancer are associated with a favorable outcome (**Fig. 3m**), since *ESR1*-positive tumors typically proliferate slower as compared to *ESR1*-negative tumors. Nonetheless, for these tumors, ERα plays a critical role in endometrial cancer development and progression^5^, the molecular actions of ERα in this cancer type remain largely unknown.

Though multi-omics analyses (**Fig. 1a**), we investigated how ERα activity is altered in endometrial cancer and how this feature may be leveraged by tumor cells to progress into the metastatic stage. Cumulatively, our findings allowed us to propose a molecular model describing the dynamic nature of ERα action in endometrial cancer development and progression (**Fig. 5i**): i) in healthy endometrial tissue, ERα regulates genes involved in cell cycle and DNA damage response (**Supplementary Fig. 1g** and **Supplementary Fig. 2c**); ii) during tumorigenesis, ERα re-localizes to the tumor-specific enhancers (**Fig. 1f-g**), including the *ESR1* enhancer itself (**Fig. 4c**). At this enhancer ERα interacts with EHMT2/G9a (**Fig. 5e**) in estradiol depend manner (**Fig. 5f**); iii) during tumor progression, the *ESR1* enhancer acquires a somatic mutation for a subset of tumors (**Fig. 4a**). *In vitro* analyses revealed that this mutation impairs EHMT2/G9a DNA binding (**Fig. 5b**), and perturbation of EHMT2/G9a in Ishikawa endometrial cancer cells enhanced *ESR1* expression (**Fig. 5g**). Given the low frequency of SNVs at ERα binding sites, many other paths to metastasis may exist, next to the mechanism we propose in this study.

Our analysis of the ERα cistrome in both normal and neoplastic endometrial tissues revealed limited overlap of chromatin occupancy between the two tissue states. Importantly, despite the healthy and tumor tissues were not matched for the same individual patient, large scale programmatic changes were consistently observed on chromatin binding of ERα, in comparing the two tissue types. Performing these analyses in matched healthy and tumor samples might lead to the identification of a higher number of significantly differentially bound sites, as noise and inter-sample variation is likely lower. This notion implies that our findings may reflect the strongest effects induced by ERα reprogramming, that are to be confirmed in larger future studies. Interestingly, tumor-depleted ERα binding sites did not display differences in H3K27ac, in contrast to the tumor-enriched ones which instead showed a consistent increase in this active histone mark. These results highlighted a specific gain-of-function for newly-engaged ERα-bound *cis*-regulatory elements in tumors, while the regulatory elements for which ERα binding is lost remain active through action of other transcription factors. Surprisingly, studying the genomic distribution of the ERα binding sites, we observed an unexpected enrichment of promoter regions for the tumor-depleted binding sites over the tumor-enriched ones. This promoter-enrichment is in contrast to prior studies, that reported ERα to primarily occupy promoter-distal *cis*-regulatory elements rather than promoters^10,14,36,37^. Importantly, as the molecular biology of this nuclear receptor has been studied mainly using cancer models, the non-tumor context of ERα action remains less well known, and may involve a higher degree of promoter involvement.

Our data, combined with previous findings showing that cell-specific ERα sites lack high-affinity estrogen responsive elements (EREs)^11^, suggest that tumor-depleted binding sites (enriched for ERα/β motifs) are essential for the cell maintenance in normal tissues. However, during tumorigenesis these sites progressively lose ERα binding in favor of sites enriched in tumors (low-affinity EREs), likely driving cell transformation. Interestingly, at these tumor-enriched binding sites we found an overrepresentation of SOX and FOX transcription factor family motifs. These finding are particularly relevant for the endometrial cancer biology since one of the most recurrently mutated genes in this cancer type is *SOX17*^28^, which acts as tumor suppressor^38^. Therefore, we can hypothesize that *SOX17* loss-of-function may leave low-affinity ERα binding sites more accessible for ERα, increasing receptor occupancy at these regions. On the other hand, enrichment of motifs for forkhead pioneer transcription factor class (FOX family) at tumor-enriched ERα binding sites is consistent with the increased H3K27ac signal at these low-affinity ERα binding sites. FOXA1 is known to facilitate ERα binding through its pioneer activity in breast cancer^39^, and we previously proposed FOXA1 to serve a comparable role in endometrial cancer^10^. In support of these hypothesis, tumor-enriched ERα binding sites display an enrichment in FOXA1 ChIP-seq peaks detected in Ishikawa endometrial cancer cells (**Fig. 1k-l** and **Supplementary Fig. 1j**). Furthermore, one can speculate on this mechanism to represent a more general feature of hormone-dependent cancers. Indeed, comparison between healthy prostate epithelium and prostate adenocarcinoma – being strongly dependent on Androgen Receptor (AR) action – showed tumor-gained AR binding sites to be selectively enriched for FOXA1 motifs, as opposed to AR sites found exclusively in healthy tissues^16^.

Enhancers interact with promoter regions to regulate gene expression, facilitated through spatial rearrangement of the chromatin structure^21^. Performing Hi-C analyses in healthy and tumor tissues, we observed a global enrichment of short-range (<2Mb) chromatin interactions, at the expense of long-range (>2Mb) contacts. Moreover, chromatin loop distance in tumor tissues is reduced at tumor-enriched ERα binding sites when compared to normal samples, while loop distance at shared and tumor-depleted sites remains unchanged. This corroborates with the observed newly acquired H3K27ac signal at tumor-enriched ERα binding sites in tumor samples in contrast to common and tumor-depleted sites, where this active enhancer/promoter histone mark is persistent in both tissue types. Combined, these findings imply a gain of enhancer activity at tumor-enriched binding sites that may lead to an increase in enhancer-promoter proximal contacts, explaining the loop distance shortening.

Next to the short/long range interaction alterations between heathy and tumor tissues, we also observed a compartment depolarization in tumor samples. Chromatin compartment destabilization is observed in many tumor types^40^ (e.g., colorectal cancer^41^ and glioblastoma^42^). Interestingly, as tumor-enriched ERα binding sites were enriched at B-to-A compartment switched sub-regions, these data suggest that alterations in 3D genome organization directly contribute to enhancer plasticity in tumorigenesis, concordant with increased H3K27ac signal at these sites.

As the vast majority of mutations are found at non-coding regions of the genome, we investigated whether tumor-enriched ERα sites could drive tumor progression through acquisition of mutations. Through integration of somatic mutation data with RNA-seq in Ishikawa cells, we concluded that most genes with proximal ERα-bound enhancer mutations are not under direct ERα control. This is in agreement with our previous observation in prostate cancer, in which only a minor fraction of SNVs at regulatory elements functionally affected their activity^17^.

We also observed a depletion of somatic mutations in tumor-depleted ERα binding sites, relative to the common and tumor-enriched ERα sites (**Fig. 3d,j**). We could speculate that two mechanisms may underlie this observation. The first is based on previous reports showing that, since DNA-bound proteins prevent efficient DNA repair^43^, somatic mutations preferentially accumulate at active TF-binding sites. On the other hand, accumulation of functional mutations may occur at tumor-enriched sites by other mechanisms. In our previous work in prostate cancer^17^, we analogously observed that tumor-gained AR binding sites were more often somatically mutated, but were also more-often positive for cancer risk SNPs. These results put our current study in perspective, suggesting that the observed higher frequency of SNVs in tumor-gained regulatory elements for hormone receptors may be a feature that is common to hormone-dependent cancers.

While the interaction between ERα and EHMT2/G9a has already been reported in breast^44,45^, EHMT2/G9a is considered to be an ERα co-activator in that cellular context. However, studies in eryhtroleukemia^46,47^ describe a dual role of EHMT2/G9a as both gene activator and repressor depending on its interaction with Mediator rather than JARID1, respectively. As we observed ERα expression levels being upregulated following knockdown of EHMT2/G9a (**Fig. 5g**), an ERα co-repressive role for EHMT2/G9a in the endometrial cancer cell context is supported by our data. Due to the lack of other ERα-positive endometrial cancer cell line availability^48^, only Ishikawa cells were used in this study. Therefore, despite experiments carried out in AN3CA ERα-negative cells support our findings, further investigation would be required to further substantiate what determines the repressive transcriptional effect of EHMT2/G9a in the interplay with ERα in this tumor type. Moreover, despite many data streams were generated in tumor tissues, EHMT2/G9a functional experiments performed in Ishikawa cells – deriving from endometrial epithelial cells – places our conclusion in an intrinsically tumor cell-centric context that ignores any potential influence of the microenvironment (e.g., stromal and immunity cells). Further studies would be required to determine whether the endometrial cancer stromal compartment may influence ERα regulation in the tumor itself.

In this study we shed light on the molecular mechanisms underlying endometrial cancer development and progression, connecting alterations in 3D genome organization with epigenetic plasticity and non-coding somatic mutations, using ESR1 enhancer mutation as an example. This study may serve as blueprint for studies in other hormone-driven cancer types, in which such complex connections are yet to be identified.

## Methods

### Human tissue collection and quality assessment

Fresh-frozen healthy and tumor samples were obtained from post-operative tissue at the Netherlands Cancer Institute (Amsterdam, the Netherlands) from patients who did not receive neoadjuvant endocrine treatment. Tumor content was assessed by haematoxylin and eosin (H&E) staining on slides taken throughout the tissue sample. For tumors, only samples displaying a tumor percentage greater or equal to 70% were deemed eligible for further analysis. Of note, non-cancerous healthy tissues were obtained from patients whose uterus was removed because of: (a) surgery for cervical carcinoma or ovarian cancer; (b) endometrial cancer diagnoses, but the area that was cryo-preserved was devoid of tumor cells.

The study was approved by the institutional review board of the Netherlands Cancer Institute, written informed consent was signed by all participants enrolled in the study, and all research was carried out in accordance with relevant guidelines and regulations.

### Cell culture and chemicals

Ishikawa (Merck Sigma Aldrich) cells were cultured in Dulbecco’s Modified Eagle Medium (DMEM, Gibco) and DMEM Mixture F-12 (DMEM/F-12, Gibco), respectively, supplemented with 10% Fetal Bovine Serum (FBS-12A, Capricorn Scientific) and penicillin/streptomycin (100μg/mL, Gibco). AN3CA endometrial cancer cells (ATCC) were cultured in Dulbecco’s Modified Eagle Medium (DMEM, Gibco) supplemented with 10% Fetal Bovine Serum (FBS-12A, Capricorn Scientific) and penicillin/streptomycin (100μg/mL, Gibco). Cell lines were subjected to regular *Mycoplasma* testing, and underwent authentication by short tandem repeat profiling (Eurofins Genomics).

For hormone stimulation, cells were pre-cultured for 3 days in phenol red-free DMEM (Gibco, Ishikawa) or DMEM/F-12 (Gibco, AN3CA) supplied with 5% dextran-coated charcoal (DCC) stripped FBS, 2mM L-glutamine (Gibco), and penicillin/streptomycin (100μg/mL, Gibco), then stimulated with 10nM DMSO-solubilized 17β-estradiol (MedChemExpress, #HY-B0141) for the indicated amount of time. Inhibition of ERα activity was performed by treatment of the cells with 100nM DMSO-solubilized Fulvestrant/ICI-182,780 (MedChemExpress, #HY-13636).

### Cell proliferation analyses

Cells were plated in a 96-well plate at a density of 10,000 cells/well. Cells were treated with 100nM DMSO-solubilized Fulvestrant/ICI-182,780 (MedChemExpress, #HY-13636). Cells were imaged every 4 hours by using an IncuCyte ZOOM Live-Cell Analysis System (Essen BioScience, Sartorious), and cell confluency percentage was calculated using the IncucyteZoom (v2018A) software. Normalization, by subtraction of the first point (T_0_), and visualization were performed using *Rseb*^49^ *(v0.3.2)* R-package (https://github.com/sebastian-gregoricchio/Rseb).

### ChIP-qPCR and ChIP-seq library preparation

Chromatin Immunoprecipitation of ERα (SantaCruz #sc-543, 5µg/IP) and H3K27ac (ActiveMotif #39133, 5µg/IP) were performed as previously described^32^ employing the combination of DSG and Formaldehyde to perform the crosslinking. The same protocol, but using 1% formaldehyde for the crosslinking, have been used to perform EHMT2/G9a (Abcam #ab133482, 5µg/IP) and Normal-IgG (Merck Millipore #12-370, same amount than IP) ChIPs in Ishikawa cells.

Immunoprecipitated DNA is compared to input sample by the comparative C_t_ method, and signal enrichment is expressed as percentage of input (% Input). Real-Time quantitative PCR (RT-qPCR) was performed on the QuantStudio™ 5 Real-Time PCR System 384-well (Applied Biosystems, #A28140) using SenSMix™ SYBR® No-ROX buffer (Meridian Bioscience). The following genomic locations have been tested: *ESR1_Enhancer1*-125kb (Fw: 5’-TGGTAGGTGCTCAGGAGATAA-3’, Rv: 5’-CAGCGACTCGAACAGGATTT-3’), *ESR1*_promoter-0.9kb (Fw: 5’-CCACTCCTGGCATTGTGATTA-3’, Rv: 5’-CAGGACACATGACACCCAAT-3’), *CDK12*_TSS (Fw: 5’-GGACCTGATCTCGCGTTGTT-3’, Rv: 5’-TAGCCTCTCGCGATGTTTCG-3’).

### ChIP-seq data processing, motif and Gene Ontology enrichment analyses

Reads were aligned to the human genome build GRCh37 using *BWA* 0.5.9-r26-dev^50^. Reads with a mapping quality (MAPQ) < 20 were removed from further analysis, and duplicates were marked using GATK markDuplicates^51^. Enrichment over input control was determined using both *MACS2*^52^ (*q* < 0.01) and *DFilter*^53^. Only peaks identified by both methods, and not overlapping with the ENCODE GRCh37/Hg19 blacklisted regions^54^, were retained. ERα consensus peaks for a given condition (healthy or normal tissues) have been defined as peaks found in at least 75% of the samples.

Differentially bound regions between tumor and healthy samples were identified using *DiffBind*^19^ with *EdgeR*^55^ mode at an FDR < 0.05. RPGC-normalized (Reads Per Genomic Content-1× coverage: (mapped reads × fragment length) / effective genome size; *bamCoverage*^56^) ChIP-seq signal at peaks was visualized using *deepTools*^56^, while genomic tracks and average density profiles were generated with *Rseb*^49^ *(v0.3.2)* R-package (https://github.com/sebastian-gregoricchio/Rseb) or *pyGenomeTracks*^57,58^.

Genomic feature annotations and distance to TSS were performed using *ChIPseeker*^59^ (promoter: −2kb:TSS:+1kb, flanking distance: 2kb), while for motif enrichment in peak regions we used *AME* v5.5.05^60^ algorithm from the *MEME* suite^61^ (v.5.5.5) using the JASPAR database^62,63^ as reference. Visualization of the enrichment motifs was made using *ggwordcloud* R-package using the *AME* computed *E*-value.

### Area proportional Venn diagrams were created using the *Vennerable* R package

Analyses for Gene Ontology biological process (GO-BP) enrichments for tumor-depleted ERα promoter-bound genes were performed using DAVID^64^ (v6.8) and employing tumor-enriched ERα promoter-bound genes as background data set. Unsupervised clustering permutation test was performed using *ConsensusClusterPlus*^65^ (v1.54.0) on the *DiffBind* normalized counts using the following parameters: reps = 100 (number of permutations), pItem = 0.8 (80% of sample shuffling), pFeature = 1 (100% of features to sample), clusterAlg = “hc” (hclust, hierarchical clustering), distance = “spearman”. To determine the ESR1/ERα motif strength at each individual ERα binding site, fasta-formatted sequences of the peaks was obtained using BED2FASTA from the *MEME* suite^61^ (v.5.5.5). Then, *PWMScore*, from *PWMTools*^66^, was used to compute the ESR1/ERα motif (JASPAR id: MA0112.3) sum occupancy score for each site. *GIGGLE*^20^ analyses were performed using the Cistrome Data Browser Toolkit (http://dbtoolkit.cistrome.org/).

### Hi-C library preparation and data processing

Flash-frozen primary tissues have been processed as described in **Fig. 2a**. Briefly, 30µm slices are cross-linked, washed, electronically homogenized and filtered using a 75µm cell strainer and collected in a 1.5mL microcentrifuge loBind tube. Then, Hi-C single-index library preparation was performed as previously described^67^ using MboI (New England Biolabs) restriction enzyme.

Quality and quantification of the Hi-C libraries was assessed using the 2100 Bioanalyzer (Agilent, DNA 7500 kit). An equimolar pool of the different samples was sequenced on the Illumina NextSeq 550 System in a 75bp paired-end setup. De-multiplexed fastq data were analyzed at 2kb, 10kb, 40kb, 100kb and 500kb resolution using the *snHiC*^68^ (v0.1.1) pipeline (https://github.com/sebastian-gregoricchio/snHiC) using default parameters and the Hg19/GRCh37 genome assembly. This pipeline relies on *HiCExplorer*^69,70^ (v3.7.2) for the matrix generation, normalization and correction, TAD and loop detection, and long/short contacts ratio and quality controls. Compartments are identified by *dcHiC*^71^ *(v2.1).* Downstream analyses have been performed using *GENOVA*^72^ (v1.0.1), which was used to plot Hi-C contact matrices heatmaps, compute Relative Contact Probabilities, and perform paired-end spatial chromatin analyses (*PE-SCAn*^23^) at ERα binding sites. *HiCExplorer*^69,70^ (v3.7.2) *hicCompartmentalization* was used to quantify the compartment polarization. For the compartment switching computation, A and B compartments locations in the reference condition were obtained merging adjacent bins displaying positive or negative, respectively, compartmentalization scores. Then, average compartmentalization score in the two conditions was computed for each reference compartment. A compartment in the reference condition was defined as “switching” when the average compartmentalization score in the reference condition switched sign in the test condition. Compartment-ChIP peaks overlaps and relative upset plot have been computed using *intervene*^73^, while Hi-C tracks were generated by *pyGenomeTracks*^57,58^.

### Whole Genome Sequencing (WGS) analyses

#### Primary samples

From the full Uterine Corpus Endometrial Carcinoma (UCEC) cohort available in the TCGA^25^ database, 41 samples have been selected in order to match the following criteria: gender = “FEMALE”, histological_type = “Endometrioid endometrial adenocarcinoma”, history_of_neoadjuvant_treatment = “No”, horm_ther = “No, I have never taken menopausal hormone therapy.”, menopause_status = “Post (prior bilateral ovariectomy OR >12 mo since LMP with no prior hysterectomy)”, prior_tamoxifen_administered_usage_category = “Never Used”, radiation_therapy = “NO”, sample_type = “Primary Tumor”, targeted_molecular_therapy = “NO”, tumor_tissue_site = “Endometrial”, *ESR1* TPM-normalized gene counts ≥ 5. Somatic mutation calling was performed on base-quality recalibrated aligned Whole Genome Sequencing data available under restricted access on the TCGA portal (data access authorization, project ID: 36269), using blood-derived matched normal samples as germline reference, by *GATK* (v 4.3.0.0) *Mutec2*^51^. Only mutations passing the GATK *FilterMutectCalls* filter and displaying a sequencing depth greater than 20 (DP>0) were retained.

#### Metastatic samples

Sampling, sequencing and initial computational steps have been previously described extensively by Priestley *et al.*^74^ and performed at the central sequencing facility at the Hartwig Medical Foundation.

Frequently mutated genes were determined by overlapping genes with somatic mutations affecting protein coding regions in more than one sample in our dataset, with the mutated genes described in the endometrial carcinoma publication from TCGA^25^ and the OncoKB Cancer Gene List^75^. Overlap between somatic variants and ChIP-seq regions was determined using *bedtools intersect* (v2.25.0)^76^.

For both datasets, mutational signatures were determined using the *deconstructSigs*^77^ R-package, using the COSMIC mutational signatures v3 database^78^. Genomic feature annotation was performed using *ChIPseeker*^59^ (promoter: −2kb:TSS:+1kb, flanking distance: 2kb).

#### Risk Single Nucleotide Polymorphism (rSNP) enrichment analyses

We assessed the enrichment of endometrial cancer germline risk variants among differentially bound ERα sites using the *VSE*^79^ (v0.99) R-package. 27 endometrial cancer risk loci (lead variant *P* < 1×10^-5^) were identified from a genome-wide association study of endometrial cancer^26^. Variants in strong LD (r^2^ > 0.8) with the lead SNP at each locus were selected using 1000 Genomes Project phase 3^80^ to generate an associated variant set (AVS). A null-distribution was built on the basis of 500 matched random variant sets. We assessed enrichment within tumor-enriched and tumor-depleted ERα binding sites, with and without including a 500bp window surrounding each binding site. A Bonferroni-corrected *P*-value < 0.05 (adjusting for four tests) was considered statistically significant.

#### Endometrial human tissue derived RNA-seq library preparation and differential expression analyses

RNA from ∼30mg of endometrial healthy or tumor tissue was extracted using the RNeasy mini kit (Qiagen) following manufacture instructions. Quality of the RNA extraction was assessed using the 2100 Bioanalyzer (Agilent, RNA 6000 Nano Kit) and selecting samples with a RNA Integrity Number (RIN) above 9. PolyA+ stranded RNA library was prepared using the Illumina Stranded mRNA Prep kit and quality was assessed by using the 2100 Bioanalyzer (Agilent, DNA 7500 kit). RNA-seq libraries have been pooled equimolarly and sequenced using NovaSeq6000 (Illumina) sequencer with a 51bp Paired-End reads setup. Fastq files have been demultiplexed by Cutadapt^81^ and mapped on Hg38/GRCh38 genome assembly using HISAT2^82^ (v2.1.0) using the following parameters: --wrapper basic-0 --min-intronlen 20 --max-intronlen 500000 --rna-strandness FR -k 5 --minins 0 --maxins 500 --fr -- new-summary. HISAT2^82^ (v2.1.0) was used to generate raw gene counts, gene counts normalization and differential expression analyses were performed using *DESeq2*^83^ (v1.30.1). Differentially expressed genes were defined by |Fold Change Expression| ≥ 2 & *P*_adj_ < 0.05. Data were visualized using *Rseb*^49^ (v0.3.2) (https://github.com/sebastian-gregoricchio/Rseb).

#### Gene Set Enrichment Analyses (GSEA)

Differentially expressed genes were ranked by the log_2_(Fold Change expression) calculated using *DESeq2*^83^ (v1.30.1) and used to perform GSEA analyses by *clusterProfiler*^84^ (v3.18.1) on the “Hallmarks” (H) datasets retrieved through *msigdbr* (v7.5.1). Visualization of the results was performed using *Rseb*^49^ (v0.3.2) (https://github.com/sebastian-gregoricchio/Rseb).

#### Public RNA-seq and differential gene expression analyses on Ishikawa cells

Ishikawa gene expression data was obtained from GEO, accession numbers <u>GSE109892</u>^30^. Differential gene expression between estradiol and DMSO treated cells was computed using *DESeq2*^83^ (v1.30.1). Differentially expressed genes were defined by |Fold Change Expression| ≥ 1.5 & P_adj_ < 0.05.

#### Circularized Chromosome Conformation Capture sequencing (4C-seq)

For each sample, 10-15 30µM-thick slices of flash frozen tissue were cross-linked under rotation at room temperature for 20 minutes in 5mL of 1× PBS supplemented with 2% FBS and 2% Methanol-free Formaldehyde. The cross-linked slices have been washed twice in 10mL of ice-cold 1× PBS before to be resuspended in 0.5-1mL of ice-cold 1× PBS in 1.5mL DNA-loBind microcentrifuge tubes. Tissues have been electronically homogenized and lysates filtered using a 75µm cell strainer. Then, experiments of 4C-seq were performed as previously described^85^. Briefly, nuclei were isolated and permeabilized to allow digestion of the chromatin by the primary RE (DpnII, New England Biolabs). Upon dilution, chromatin fragments have been ligated and then de-crosslinked. DNA was purified and digested with the secondary RE (NlaIII, New England Biolabs) and circularized by ligation. PCR amplification of the re-purified circular fragments have been performed using View-Point specific primers (Reading Primer: 5’-tacacgacgctcttccgatctAACTCGATTTGGAGCGATC-3’; Non-reading Primer: 5’-actggagttcagacgtgtgctcttccgatctCTGGGACTGCACTTGCTC-3’) using the Expand Long Template PCR System (Roche). Amplicons were purified with AMPure XP beads (Beckman Coulter) in a 0.8× ratio and then amplified using standard indexed Illumina primers as previously described^85^ using the Expand Long Template PCR System (Roche). Second-round PCR products were purified with PCR purification columns (Qiagen) and quantified by 2100 Bioanalyzer (Agilent, DNA 7500 kit).

4C libraries have been pooled equimolarly and sequenced using Illumina MiSeq with a 75bp Single-End reads setup. Fastq files have been demultiplexed by Cutadapt^81^ and mapped on Hg19/GRCh37 genome assembly and signal normalized by *pipe4C*^85^ v1.1 R-package in “cis” mode and with default parameters. The signal of different technical replicates was averaged to obtain a unique signal for each sample. The, each tissue sample have been considered as biological replicate. Downstream analyses of 4C-seq data have been performed in an R v4.0.3 environment by using *get.single.base.score.bw* and *genomic.track* functions from *Rseb*^49^ (v0.3.2) (https://github.com/sebastian-gregoricchio/Rseb) R-package in combination with *ggplot2* (v3.3.5) and *ggforce* (v0.3.3) R-packages.

#### Protein extraction and immunoblotting

For immunoprecipitated proteins, elution buffer (1% SDS, 0.1M NaHCO_3_) and Laemmli buffer, supplied with 100mM DTT (Dithiothreitol, Merck Sigma Aldrich), were added to input samples and beads, and then boiled at 95°C for 30min. For whole-cell extracts, cells have been lysed using 2× Laemmli buffer supplied with 100mM DTT (Dithiothreitol, Merck Sigma Aldrich), boiled at 95°C for 10min and then DNA was sheared by sonication (Bioruptor® Pico, Diagenode; 4 cycles of 30s ON + 90s OFF). In both cases, proteins were then resolved by SDS-PAGE and transferred to a 0.22μm nitrocellulose membrane (BioRad). Membranes were incubated for 2h with blocking solution (BS: 1× PBS, 0.1% Tween20, 5% non-fat milk powder) and then incubated with primary antibody against ERα (ThermoFisher #MA5-14104, 1:1000 in BS), EHMT2/G9a (Abcam #ab133482, 1:1000 in BS) or GAPDH (ThermoFisher #PA1-987, 1:5000 in BS) overnight at +4°C. After washing with washing solution (1× PBS, 0.1% Tween20), membranes were incubated with secondary antibodies donkey-α-mouse IRDye® 680RD (926-68073, LI-COR Biosciences, 1:10,000 in BS), donkey-α-mouse IRDye® 800CW (926-32212, LI-COR Biosciences, 1:10,000 in BS), donkey-α-rabbit IRDye® 800CW (926-32213, LI-COR Biosciences, 1:10,000 in BS) or donkey-α-rabbit IRDye® 680RD (926-68073, LI-COR Biosciences, 1:10,000 in BS) for 1h. Membranes were washed again with washing solution, scanned and analyzed using an Odyssey® CLx Imaging System (LI-COR Biosciences) and *ImageStudio™ Lite* v.5.2.5 software (LI-COR Biosciences).

#### Sequence-based Machine Learning 3D genome folding predictions

The 3D genome organization perturbation predictions have been performed using a window of 2^20^bp around the *ESR1* locus employing the Convolutional Neural Networks (CNN) model from *Akita*^34^ tool. We used the pre-existing model based on Hi-C data originated from human foreskin fibroblast (HFF). Heatmaps have been generated using *matplotlib.pyplot.matshow*.

#### DNA-oligo protein pull-down

Ishikawa cells were harvested and washed twice with ice-cold PBS and nuclear extracts were prepared as described previously^86^. Briefly, cells are washed with 1× PBS and trypsinized. Trypsin is neutralized by adding the appropriate SILAC medium, and “light”/”heavy” cells are collected separately at 4°C. Cells are washed in 1× ice-cold PBS and resuspended in five volumes of ice-cold buffer A (10 mM KCl, 20 mM Hepes KOH pH 7.9, 1.5 mM MgCl_2_). Cells are then pelleted and resuspended in two volumes of buffer A supplemented with protease inhibitors and 0.15% Igepal NP40 (v/v) before to be dounce homogenized. Nuclei are then collected and resuspended in two volumes of buffer C (420 mM NaCl, 20 mM Hepes KOH pH 7.9, 20% glycerol (v/v), 2 mM MgCl_2_, 0.2 mM EDTA, and 0.1% Igepal CA630 (NP40; Sigma-Aldrich, I8896), EDTA-free complete protease inhibitors (Roche), and 0.5 mM DTT) and incubated for 1 hour. The suspension is centrifuged, and the supernatant containing the nuclear extracts is collected.

The ∼50bp oligonucleotide probes encompassing the SNP were ordered with the forward strand containing a 5’-biotin moiety (Integrated DNA Technologies) (Fw_WT: 5’-/5Biosg/ACTGTGGAAACTGGAAGCTG<u>TTC</u>TTGGACTATTTCGCAACACTTTTCTCC-3’; Rv_WT: 5’-GGAGAAAAGTGTTGCGAAATAGTCCAAGAACAGCTTCCAGTTTCCACAGT-3’; Fw_SNP: 5’-/5Biosg/ACTGTGGAAACTGGAAGCTG<u>T</u>TTGGACTATTTCGCAACACTTTTCTCCTC-3’, Rv_SNP: (5’-GAGGAGAAAAGTGTTGCGAAATAGTCCAAACAGCTTCCAGTTTCCACAGT-3’; mutated site is underlined). DNA affinity purifications, on-bead trypsin digestion and dimethyl labeling were performed as described^87^. Matching light and medium labelled samples were then combined and analyzed using a gradient from 7 to 30% buffer B in Buffer A over 44min, followed by a further increase to 95% in the next 16min at flow rate of 250nL/min using an Easy-nLC 1000 (Thermo Fisher Scientific) coupled online to an Orbitrap Exploris 480 (Thermo Fisher Scientific). MS1 spectra were acquired at 120,000 resolution with a scan range from 350 to 1300Lm/z, normalized AGC target of 300% and maximum injection time of 20 ms. The top 20 most intense ions with a charge state 2-6 from each MS1 scan were selected for fragmentation by HCD. MS2 resolution was set at 15,000 with a normalized AGC target of 75%. Raw MS spectra were analyzed using MaxQuant software (version 1.6.0.1) with standard settings^87,88^. Data was searched against the human UniProt database (downloaded 2017) using the integrated search engine. N-terminal and lysine modification for dimethyl labeling was specified under “labels”. Carbamidomethylation was specified as a fixed modification. N-terminal acetylation and methionine oxidation were selected as variable modifications.

#### Transient small interfering RNA (siRNA) cell transfection

Transient transfections of Ishikawa and AN3CA cell lines was performed according to the manufacturer’s instructions using Lipofectamine™ RNAiMAX (Invitrogen) for siRNA knockdown experiments. The ON-TARGETplus™ siRNA SMARTpool targeting human EHMT2/G9a (L-006937-00-00050) and the siGENOME™ Non-Targeting control siRNA #5 (D-001210-05-20) were purchased from Dharmacon.

#### Statistical analysis

All statistical analyses were performed in R versions 3.4.4, 3.5.1 or 4.0.2 (R Core Team 2020, https://www.R-project.org). Enrichment of Ishikawa peaks and somatic mutations in differentially bound regions was calculated using Fisher’s exact test, and, where appropriate, corrected for multiple testing using the FDR.

## Data availability

Endometrial healthy and tumor tissues ERα and H3K27ac ChIP-seq, RNA-seq, Hi-C and, 4C-seq raw sequencing data (GRCh37/Hg19 genome build) have been deposited at the European Genome-phenome Archive (EGA), which is hosted by the EBI and the CRG, under accession number EGAS00001007240, while endometrial tissue processed data and Ishikawa ChIP-seq data are deposited at the Gene Expression Omnibus (GEO) database under accession number GSE235241.

Raw sequencing data of the RNA-seq performed in Ishikawa cells can be found at GEO database under accession number GSE109892^30^. ERα ChIP-seq data in breast cancer tissues from Singh *et al.*^32^ are accessible in GEO under accession number <u>GSE114737</u>. Ishikawa ChIP-seq data in the form of optimal Irreproducibility Discovery Rate (IDR) thresholded peaks (GRCh37/Hg19 genome build) were obtained from the ENCODE data portal^48^ (**Supplementary Table 2**). The H3K27ac HiChIP data in Ishikawa cells have been obtained from O’Mara *et al.*^89^ accessible in GEO under accession number <u>GSE137936</u>. Mass spectrometry data have been deposited at the ProteomeXchange Consortium through the PRIDE^90^ partner repository with the identifier <u>PXD029822</u>. The breast and endometrial cancer ATAC-seq, gene expression, progression-free survival and Whole Genome Sequencing (data access authorization for project ID: 36269) used in this this work are in whole or part based upon data generated by the TCGA Research Network^25,31^ (https://www.cancer.gov/tcga).

## Acknowledgements

We thank members of the Zwart and Bergman labs for valuable feedback, suggestions and input. We would like to acknowledge the Research High Performance Computing (RHPC) facility of the Netherlands Cancer Institute (NKI) to have enabled us to perform all the computations required to analyze the data generated, the NKI Genomics Core Facility for next-generation sequencing and bioinformatics support, and the NKI-AVL Core Facility Molecular Pathology and Biobanking for patient samples handling. This publication and the underlying study have been made possible partly based on data that Hartwig Medical Foundation has made available to the study through the Hartwig Medical Database. Further, we would like to thank Hartwig Medical Foundation for facilitating and supporting whole-genome sequencing data generation, data analyses and data interpretation.

## Author contributions

Conceptualization: S.G., A.K and W.Z. W.Z. was responsible for project funding. S.G. and A.K. performed Hi-C and 4C-seq experiments; S.G., K.S. and M.D. performed ChIP/ChIP-seq and immunoblotting experiments; S.S. performed and analyzed the protein DNA-oligo pulldowns experiments; S.G., M.H, T.M.S. and A.A.S. analyzed the omics data; T.A.O. and D.M.G. provided support in the HiChIP and rSNP data analyses; M.V. provided support for the proteomics experiments; L.F.A.W. provided bioinformatics support; F.v.L. designed and performed study for patient sample collection. S.G., A.K. and W.Z. wrote the manuscript, with input from all co-authors.

## Supplementary Figures

**Supplementary Fig. 1:** Endometrial cancer tissues and ChIP-seq quality controls. **a** Hematoxylin and eosin stain of healthy (top) and tumor (bottom) representative endometrial tissues. These tissues correspond to tissues used for Hi-C and RNA-seq analyses. **b** Bar plot (top) and box plot (bottom) showing the total number of reads of the ERα (black) and H3K27ac (red) ChIP-seq libraries among all the healthy (blue) and tumor (orange) tissue samples. Wilcoxon test: *P*_ERα_ = 0.659, *P*_H3K27ac_ = 0.21 (*ns* = not significant, *P*≥0.05). **c** Principal Component Analyses (PCA) of ERα (circles) and H3K27ac (triangles) ChIP-seq data in 4 healthy (blues) and 5 tumor (orange) endometrial primary tissues. **d** Unsupervised permutation clustering test results. Gradient indicates the clustering probability between samples. **e** Distribution of the number of ERα differential peaks obtained by 100 iterations of label reshuffling. **f** Boxplot of the normalized read count distribution at ERα bindings sites detected in healthy (blue) or tumor (orange) tissues, as well as at differential consensus ERα bindings sites with lower (–) or higher (+) affinity in tumor tissues. **g** Bubble plot showing the relative enrichment of DAVID Gene Ontology biological processes (GO-BP) enrichments for tumor-depleted ERα promoter-bound genes. Tumor-enriched ERα promoter-bound genes were used as background data set. **h** Stacked bar plot showing the presence (PWMScore > 0) frequency of ESR1/ERα motif in different ERα binding categories. **i** Bar plot depicting the average PWMScore ± SEM at different ERα binding categories. *P* value of Wilcoxon test is indicated. **j** Top 20 ranked factors identified to be enriched at different ERα binding categories as identified by GIGGLE^20^ analyses.

**Supplementary Fig. 2:** Differential gene expression between normal and tumor endometrial tissues. **a** Principal Components analyses of RNA-seq data for 3 healthy (blue shades) and 3 tumor (orange shades) samples. Scatter plot between PC1 and PC2 is shown. **b** Volcano plot of the differential gene expression analyses between 3 normal (blue) and 3 tumor (orange) endometrial tissues. Vertical lines indicate a Fold Change Expression equal to 2 (positive side)/0.5 (negative side), while the horizontal line indicates a *P*_adj_ = 0.05. **c** Violin plot showing the *DESeq2* normalized counts in healthy (blue) vs tumor (orange) tissues for the genes included in the over-represented biological processes identified in **Supplementary** Fig. 1g. *P*-value of a paired Wilcoxon test is indicated above each category, in gray when above 0.05, and in black when below 0.05 (statistically significant). **d** GSEA analyses on full transcriptome based on genes ranked by fold change differential expression between tumor and normal tissues. Bar plot of the Normalized Enrichment Score (NES) is shown for statistically significant gene sets (*P* < 0.05). **e** Stacked plot of the differential expression frequency of genes associated to different classes of ERα binding sites. **f** Volcano plot of the differential gene expression analyses between 3 normal (blue) and 3 tumor (orange) endometrial tissues of genes associated with different classes of ERα binding sites. Vertical lines indicate a Fold Change Expression equal to 2 (positive side)/0.5 (negative side), while the horizontal line indicates a *P*_adj_ = 0.05.

**Supplementary Fig. 3:** 3D genome organization analyses in endometrial primary tissues. **a** Chromosome matrix for each individual sample showing the chromosome translocation score per each chromosome based on the Hi-C contact probability (500kb resolution). **b** Pearson Hi-C contacts correlation matrix among different healthy (blue) and tumor (orange) endometrial primary tissues. **c** Stacked bar plot showing the Hi-C contact type frequency per each sample, as computed by *HiCExplorer*. Numbers indicate the millions of Hi-C contacts (paired-end fragments) detected. **d** Average Relative Hi-C Contact Probability as function of the distance for healthy and tumor samples (40kb resolution). **e** Violin plot of the average distribution of the short-range (<2Mb) over long-rage (>2Mb) Hi-C contacts ratio in healthy and tumor samples (40kb resolution). Wilcoxon’s test *P*-value is indicated. **f** Average compartment polarization ratio (100kb resolution), defined as (AA + BB) / (AB + BA), for healthy and tumor samples. **g** On the upper part, average saddle plot of A/B compartments interactions as computed in (**f**) for each tissue type. On the lower part, Tumor-Healthy difference of saddle-score, where orange indicates a higher score in tumor samples while purple a higher score in healthy tissues. **h** On the left, scatter plot showing the correlation of average individual compartment bin scores in tumor vs healthy endometrial tissues and overlapping with the different categories of ERα consensus peaks. On the right, bin count fold change for each compartment transition subcategory (A-to-A, B-to-B, A-to-B, B-to-A) relative to the count of bins not overlapping with any ERα binding site. **i** Quantification of the average 4C-seq signal from Fig. 4c at the “*Enhancer 1*” and “*Enhancer 2*” regions per each tissue sample. Each point represents the replicate mean value of each sample. Genomic location (GRCh37/Hg19) of the two regions are indicated in parenthesis.

**Supplementary Fig. 4:** Example of chromatin loops in healthy and tumor tissues Heatmaps depict the averaged 10kb resolution Hi-C matrix for the 3 healthy (upper) and 3 tumor (lower) tissues. Loops specific to healthy (blue) or tumor (orange) tissues are indicated by squares on the heatmaps, while the dashed lines represent the diagonal projection of the loop anchors. Loops are indicated by arcs that connect the loop anchors. Genomic tracks are showing the normalized ChIP-seq signal for ERα (top tracks) and H3K27ac (bottom tracks) in healthy (blue) and tumor (orange) endometrial tissues.

**Supplementary Fig. 5:** *ESR1* gene expression in normal and tumor endometrial human samples **a** RNA-seq *ESR1* gene expressed as *DESeq2* normalized counts between healthy and tumor endometrial tissues from data generated in this work. Each individual sample is represented by a dot. *P* value of Wilcoxon test is indicated. **b** Violin and box plot showing the RNA-seq *ESR1* gene expression from endometrial carcinoma TNMplot data using normal samples from non-cancerous patients as control. Average of 146 healthy and 547 tumor samples is shown. *P* value of Wilcoxon test is indicated. **c** Violin and box plot showing the RNA-seq *ESR1* gene expression from endometrial carcinoma TNMplot data using paired tumor and adjacent normal tissues samples. Average of 23 patients is shown. *P* value of paired Wilcoxon test is indicated.

**Supplementary Fig. 6:** Somatic mutation overlaps with ER**_α_** binding sites in hypermutated and non-hypermutated primary and metastatic samples **a-b** Stacked bar blot depicting the fraction of ERα peaks overlapping with somatic mutations identified in the primary (**a**) or metastatic (**b**) patient cohort. Overlaps of mutation occurring in non-hypermutated (*left*) and hypermutated (*right*) are shown independently.

**Supplementary Fig. 7:** Genomic and epigenomic landscape comparison of the *ESR1* locus in endometrial and breast cancer. **a** Source images and Ponceau Red staining of the western blot shown in Fig. 3l. Cropped areas in the main figure are indicated by a red rectangle. **b** Average density profile of ATAC-seq signal from publicly available data (TCGA) for endometrial and breast cancer at the *ESR1* locus (*Enhancer 1*, *Enhancer 2* and promoter). Signal was smoothed using the loess regression method. **c** ChIP-seq genomic tracks for ERα in healthy endometrial tissues (top, blue), endometrial tumors (middle, orange) and breast cancer (bottom, pink) at the *ESR1* locus. Focused average density signal at *ESR1 Enhancer 1*, *Enhancer 2* and promoter are plotted on the lower panel. Average density signal was smoothed using the loess regression method.

**Supplementary Fig. 8:** Validation of the TCGA gene expression analyses. **a** Density distribution of the *ESR1* gene expression correlation with the whole transcriptome in TCGA RNA-seq data. Vertical blue and red lines indicate the gene expression correlation score between *ESR1* and the top 3 anti-correlated and 3 correlated genes showed in Fig. 5c. Black dotted vertical lines indicate respectively the 5^th^, 50^th^ (median) and 95^th^ percentile of the distribution. **b** Progression-Free Kaplan-Meier curve of endometrial cancer patients (TCGA data) divided into two groups using the median of expression gene (FPKM) as cut-off: blue high expression, red low expression. On the upper part are shown the curves for *GTF2IRD1*, *ZNF768*, *SIN3A*, while on the lower part *XPA*, *ZBTB21*, *POLK*. **c-d** Source images and Ponceau Red staining of the western blot shown in Fig. 5e (**c**) and Fig. 5g (**d**). Cropped areas in the main figure are indicated by red rectangles.

**Supplementary Fig. 9:** EHMT2/G9a functional analyses in AN3CA ER**_α_**-deficient endometrial cancer cell lines **a** Whole protein extracts from Ishikawa and AN3CA endometrial cancer cell lines were immunoblotted using antibodies against EHMT2/G9a, ERα and GAPDH (loading control). **b** Source images and Ponceau Red staining of the western blot shown in (**a**). Cropped areas in the main figure are indicated by red rectangles. **c-d** Normalized cell confluency of Ishikawa (**c**) and AN3CA (**d**) endometrial cancer cells upon DMSO (black) or 100nM ICI-182,780/Fulvestrant (green) treatment over time. Average ± SEM of 6 replicates is shown. **e** EHMT2/G9a ChIP in AN3CA cells stimulated for 6h with 10nM β-estradiol. Bar plot shows percentage of enrichment over the input (% Input) at the *ESR1 Enhancer 1*, *ESR1* promoter and *CDK12* promoter (negative control) analyzed by quantitative PCR (qPCR). Mean of 2 independent experiments is shown. **f** AN3CA endometrial cancer cells were stimulated for 72h with 10nM β-estradiol in combination or not with 100nM ICI-182,780 (Fulvestrant). In all conditions, cells where incubated either with a Non-Targeting (NT) siRNA or with an siRNA against *EHMT2/G9a*. Then, whole-cell extracts were analyzed by immunoblotting with antibodies against ERα, EHMT2/G9a and GAPDH (loading control). **g** Source images and Ponceau Red staining of the western blot shown in (**f**). Cropped areas in the main figure are indicated by red rectangles.

## Table legends

**Supplementary Table 1:** Clinicopatholigical features of patient samples and omics data streams generated. Description of the collection site, histological status, site of collection, age at diagnosis, tumor percentage, grade and FIGO classification for each patient sample. The last column indicates the omics data generated.

**Supplementary Table 2:** List of publicly available ChIP-seq data sets in Ishikawa cells. List of transcription and epigenetic factors ChIP-seq data in Ishikawa cells and corresponding ENCODE accession number.

**Supplementary Table 3:** Metadata relative to metastatic endometrial cancer samples. In this tables are reported the fraction of tumor cells, year of birth, biopsy site and eventual pre-treatments of the 24 metastatic endometrial cancer samples.

**Supplementary Table 4:** List of ER**_α_** binding sites and H3K27ac HiChIP linked target genes List of candidate target genes for the tumor-gained mutated ERα sites based on H3K27ac HiChIP analysis of Ishikawa cells^89^. The structure of the table is the following: *chr_enh/start_enh/end_enh*: genomic position of the ERα binding sites; *count_enh*: enhancer identifier; *target_gene/ensemble_ID*: candidate target gene of enhancer from endometrial cancer H3K27ac HiChIP data; *enhancer_in_HiChIP_target_gene_promoter_anchor?*: *yes*, enhancer is located within the HiChIP anchor that encompasses the gene TSS – *no*, enhancer is located in the HiChIP anchor distal to the gene; *enhancer_ within_3kb_TSS_of_target_gene?*: *yes*, enhancer is located within ±3kb of the TSS of the gene – *no*, enhancer is not located within 3kb of the TSS of the gene.

