## Supplementary Fig. for "Enhancer plasticity in endometrial tumorigenesis demarcates non-coding somatic mutations and 3D-genome alterations boosting the oncogenic driver ESR1"

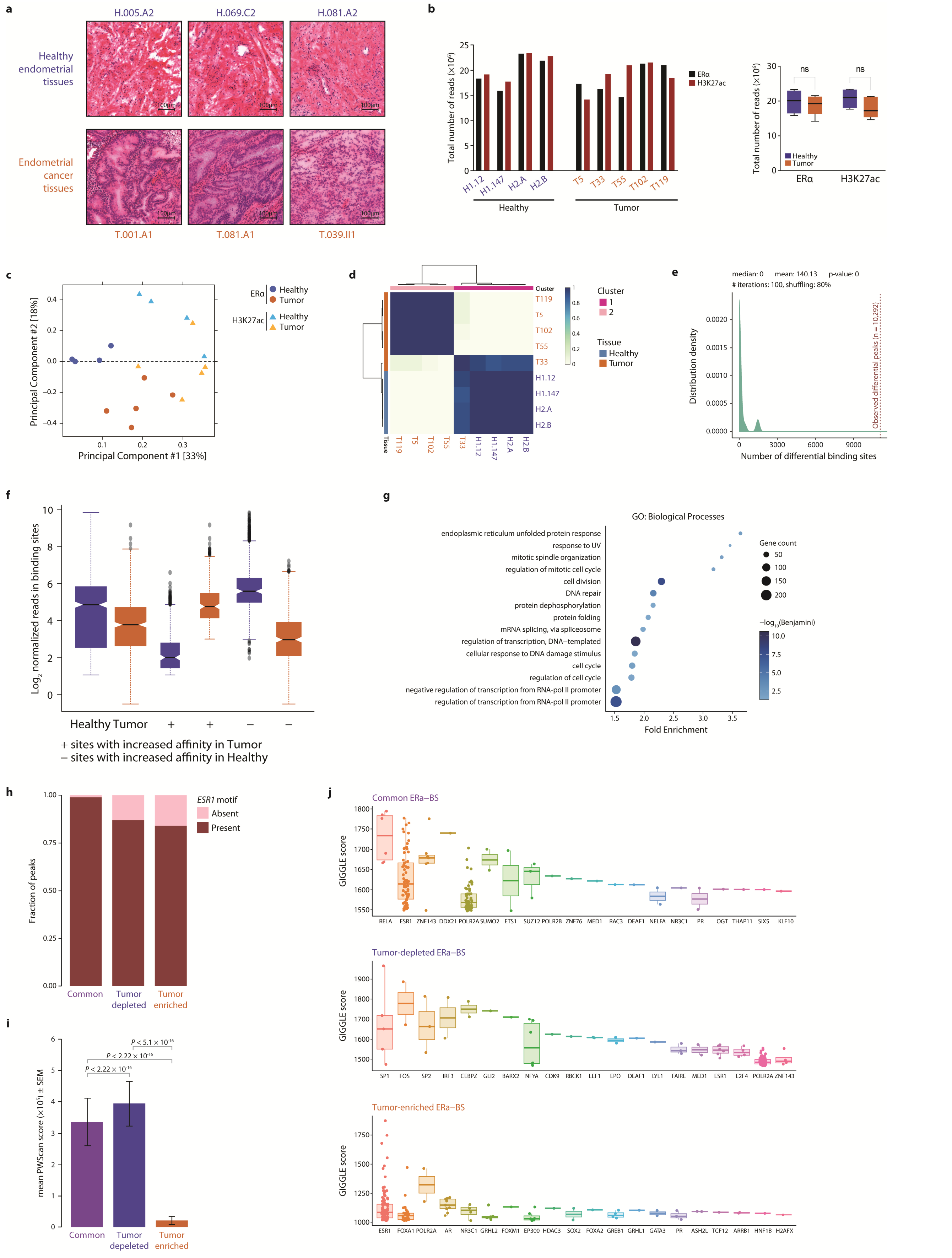


**Supplementary Fig. 1: Endometrial cancer tissues and ChIP-seq quality controls.**

**Supplementary Fig. 1: Endometrial cancer tissues and ChIP-seq quality controls.**

**a** Hematoxylin and eosin stain of healthy (top) and tumor (bottom) representative endometrial tissues. These tissues correspond to tissues used for Hi-C and RNA-seq analyses. **b** Bar plot (top) and box plot (bottom) showing the total number of reads of the ERα (black) and H3K27ac (red) ChIP-seq libraries among all the healthy (blue) and tumor (orange) tissue samples. Wilcoxon test: $P_{ER\alpha} = 0.659$, $P_{H3K27ac} = 0.21$ (ns = not significant, P≥0.05). **c** Principal Component Analyses (PCA) of ERα (circles) and H3K27ac (triangles) ChIP-seq data in 4 healthy (blues) and 5 tumor (orange) endometrial primary tissues. **d** Unsupervised permutation clustering test results. Gradient indicates the clustering probability between samples. **e** Distribution of the number of ERα differential peaks obtained by 100 iterations of label reshuffling. **f** Boxplot of the normalized read count distribution at ERα bindings sites detected in healthy (blue) or tumor (orange) tissues, as well as at differential consensus ERα bindings sites with lower (–) or higher (+) affinity in tumor tissues. **g** Bubble plot showing the relative enrichment of DAVID Gene Ontology biological processes (GO-BP) enrichments for tumor-depleted ERα promoter-bound genes. Tumor-enriched ERα promoter-bound genes were used as background data set. **h** Stacked bar plot showing the presence (PWMScore > 0) frequency of ESR1/ERα motif in different ERα binding categories. **i** Bar plot depicting the average PWMScore ± SEM at different ERα binding categories. *P* value of Wilcoxon test is indicated. **j** Top 20 ranked factors identified to be enriched at different ERα binding categories as identified by GIGGLE analyses.

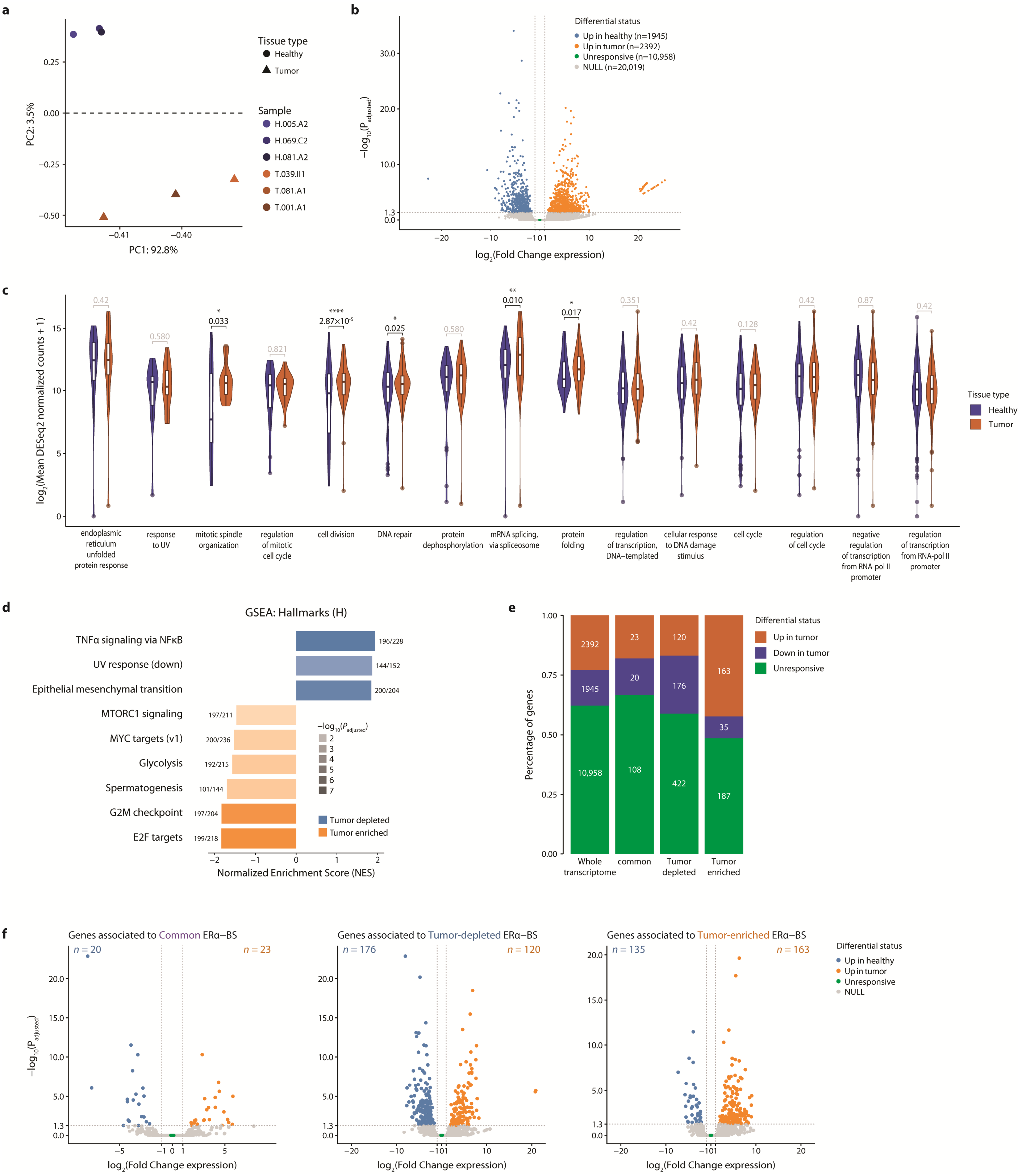


**Supplementary Fig. 2: Differential gene expression between normal and tumor endometrial tissues.**

**Supplementary Fig. 2: Differential gene expression between normal and tumor endometrial tissues.**

**a** Principal Components analyses of RNA-seq data for 3 healthy (blue shades) and 3 tumor (orange shades) samples. Scatter plot between PC1 and PC2 is shown. **b** Volcano plot of the differential gene expression analyses between 3 normal (blue) and 3 tumor (orange) endometrial tissues. Vertical lines indicate a Fold Change Expression equal to 2 (positive side)/0.5 (negative side), while the horizontal line indicates a $P_{adj} = 0.05$. **c** Violin plot showing the *DESeq2* normalized counts in healthy (blue) vs tumor (orange) tissues for the genes included in the over-represented biological processes identified in Supplementary Fig. 1g. *P*-value of a paired Wilcoxon test is indicated above each category, in gray when above 0.05, and in black when below 0.05 (statistically significant). **d** GSEA analyses on full transcriptome based on genes ranked by fold change differential expression between tumor and normal tissues. Bar plot of the Normalized Enrichment Score (NES) is shown for statistically significant gene sets ($P < 0.05$). **e** Stacked plot of the differential expression frequency of genes associated to different classes of ERα binding sites. **f** Volcano plot of the differential gene expression analyses between 3 normal (blue) and 3 tumor (orange) endometrial tissues of genes associated with different classes of ERα binding sites. Vertical lines indicate a Fold Change Expression equal to 2 (positive side)/0.5 (negative side), while the horizontal line indicates a $P_{adj} = 0.05$.

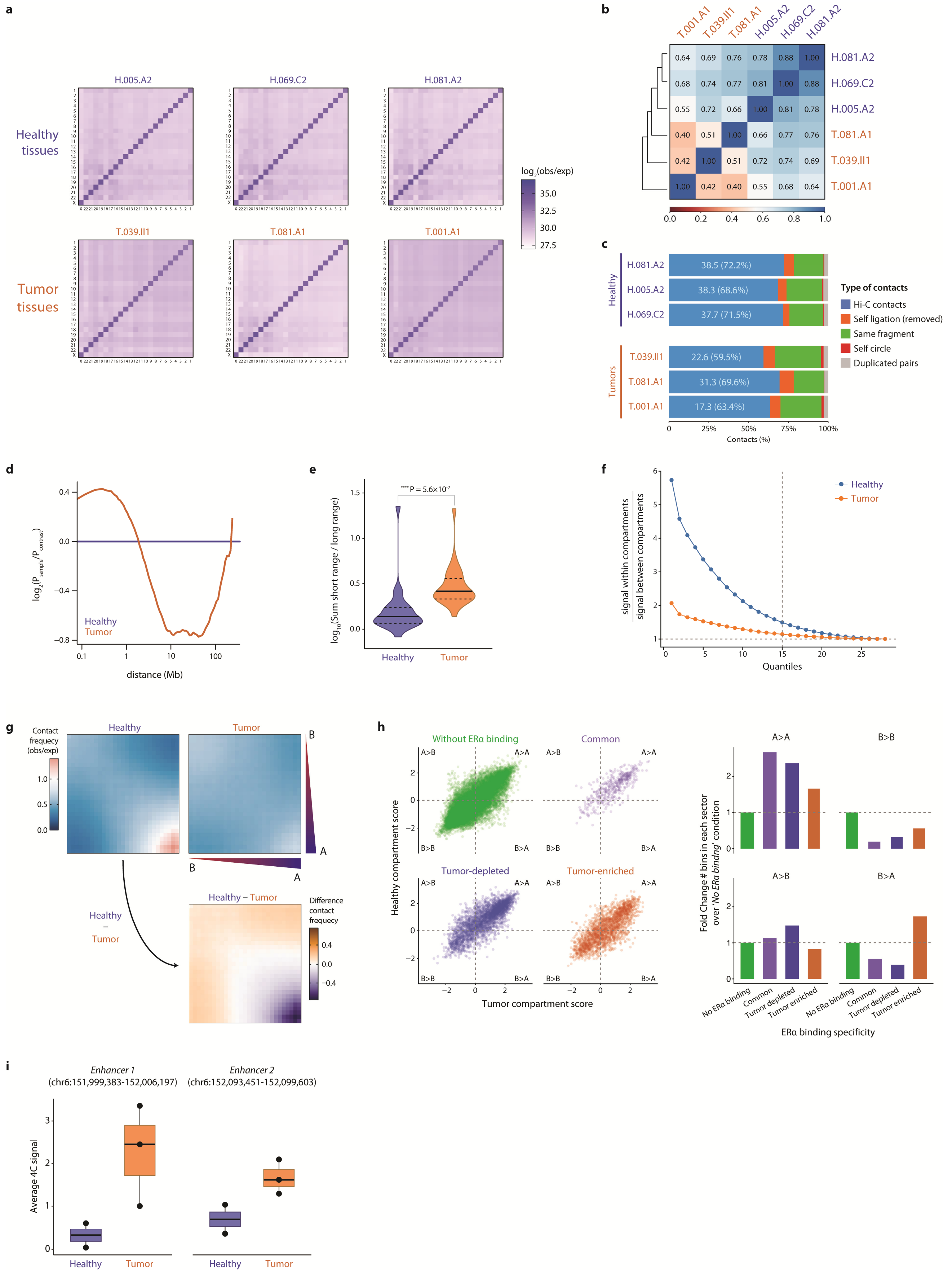


**Supplementary Fig. 3: 3D genome organization analyses in endometrial primary tissues.**

**Supplementary Fig. 3: 3D genome organization analyses in endometrial primary tissues.**

**a** Chromosome matrix for each individual sample showing the chromosome translocation score per each chromosome based on the Hi-C contact probability (500kb resolution). **b** Pearson Hi-C contacts correlation matrix among different healthy (blue) and tumor (orange) endometrial primary tissues. **c** Stacked bar plot showing the Hi-C contact type frequency per each sample, as computed by *HiCExplorer*. Numbers indicate the millions of Hi-C contacts (paired-end fragments) detected. **d** Average Relative Hi-C Contact Probability as function of the distance for healthy and tumor samples (40kb resolution). **e** Violin plot of the average distribution of the short-range (<2Mb) over long-rage (>2Mb) Hi-C contacts ratio in healthy and tumor samples (40kb resolution). Wilcoxon's test *P*-value is indicated. **f** Average compartment polarization ratio (100kb resolution), defined as (AA + BB) / (AB + BA), for healthy and tumor samples. **g** On the upper part, average saddle plot of A/B compartments interactions as computed in (**f**) for each tissue type. On the lower part, Tumor-Healthy difference of saddle-score, where orange indicates a higher score in tumor samples while purple a higher score in healthy tissues. **h** On the left, scatter plot showing the correlation of average individual compartment bin scores in tumor vs healthy endometrial tissues and overlapping with the different categories of ERα consensus peaks. On the right, bin count fold change for each compartment transition subcategory (A-to-A, B-to-B, A-to-B, B-to-A) relative to the count of bins not overlapping with any ERα binding site. **i** Quantification of the average 4C-seq signal from **Fig. 4c** at the "*Enhancer 1*" and "*Enhancer 2*" regions per each tissue sample. Each point represents the replicate mean value of each sample. Genomic location (GRCh37/Hg19) of the two regions are indicated in parenthesis.

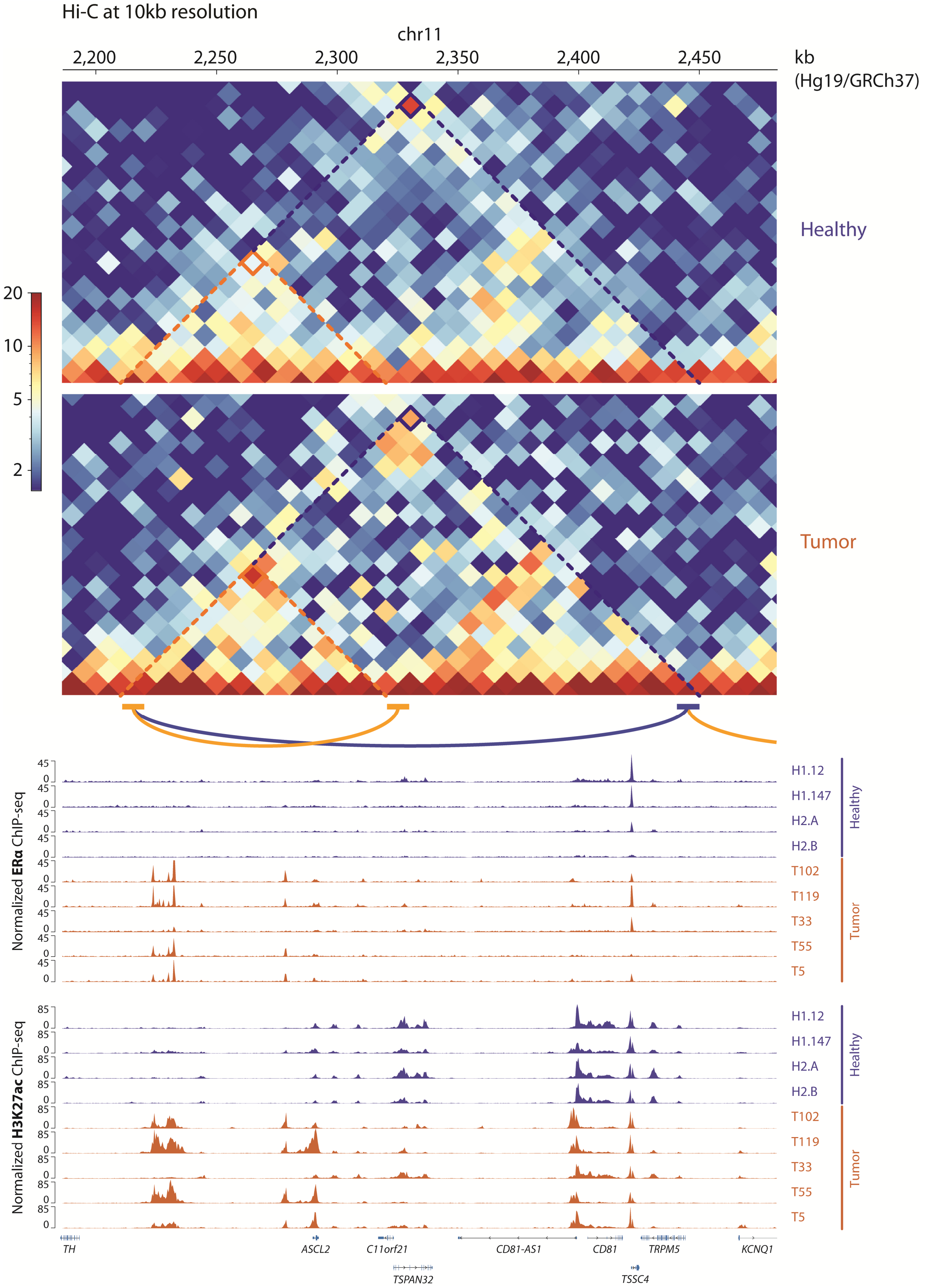


**Supplementary Fig. 4: Example of chromatin loops in healthy and tumor tissues**

**Supplementary Fig. 4: Example of chromatin loops in healthy and tumor tissues**

Heatmaps depict the averaged 10kb resolution Hi-C matrix for the 3 healthy (upper) and 3 tumor (lower) tissues. Loops specific to healthy (blue) or tumor (orange) tissues are indicated by squares on the heatmaps, while the dashed lines represent the diagonal projection of the loop anchors. Loops are indicated by arcs that connect the loop anchors. Genomic tracks are showing the normalized ChIP-seq signal for ERα (top tracks) and H3K27ac (bottom tracks) in healthy (blue) and tumor (orange) endometrial tissues.

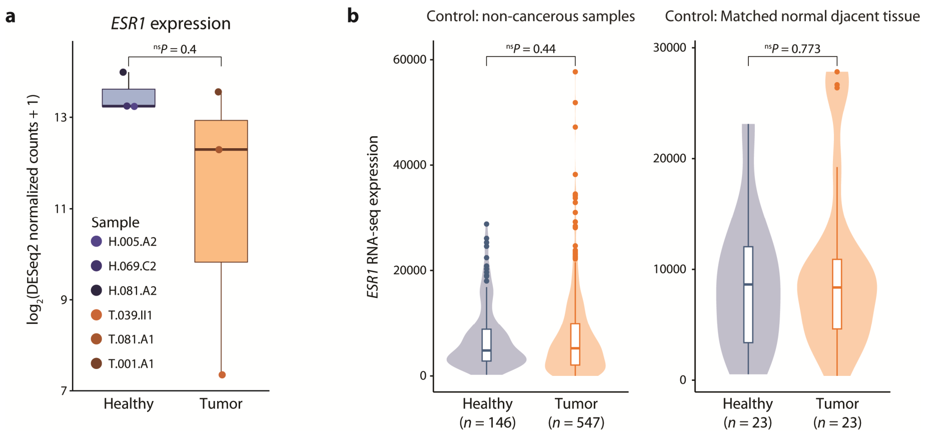


### Supplementary Fig. 5: *ESR1* gene expression in normal and tumor endometrial human samples

**a** RNA-seq *ESR1* gene expressed as *DESeq2* normalized counts between healthy and tumor endometrial tissues from data generated in this work. Each individual sample is represented by a dot. *P* value of Wilcoxon test is indicated. **b** Violin and box plot showing the RNA-seq *ESR1* gene expression from endometrial carcinoma TNMplot data using normal samples from non-cancerous patients as control. Average of 146 healthy and 547 tumor samples is shown. *P* value of Wilcoxon test is indicated. **c** Violin and box plot showing the RNA-seq *ESR1* gene expression from endometrial carcinoma TNMplot data using paired tumor and adjacent normal tissues samples. Average of 23 patients is shown. *P* value of paired Wilcoxon test is indicated.

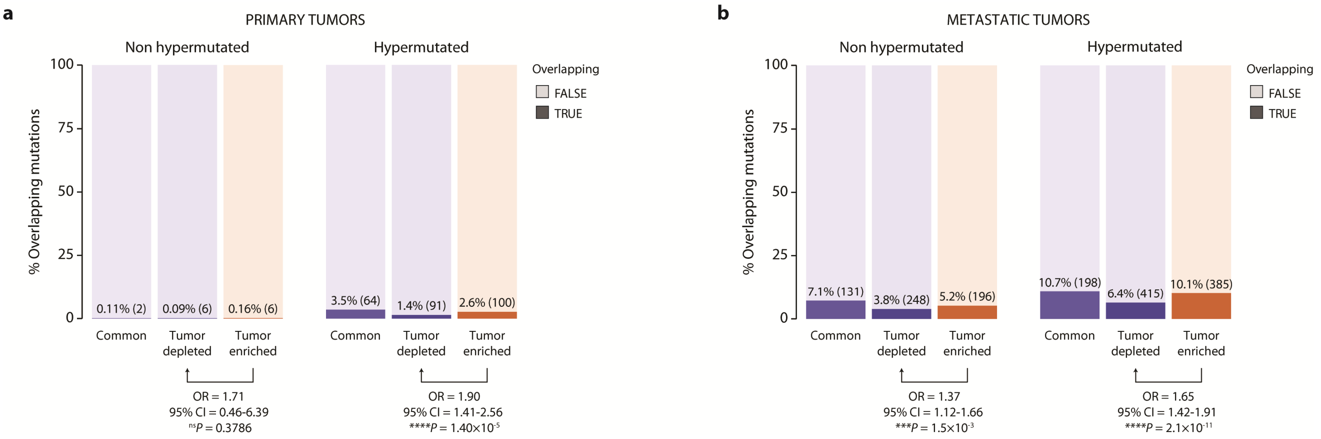


### Supplementary Fig. 6: Somatic mutation overlaps with ERα binding sites in hypermutated and non-hypermutated primary and metastatic samples

**a-b** Stacked bar blot depicting the fraction of ERα peaks overlapping with somatic mutations identified in the primary (**a**) or metastatic (**b**) patient cohort. Overlaps of mutation occurring in non-hypermutated (*left*) and hypermutated (*right*) are shown independently.

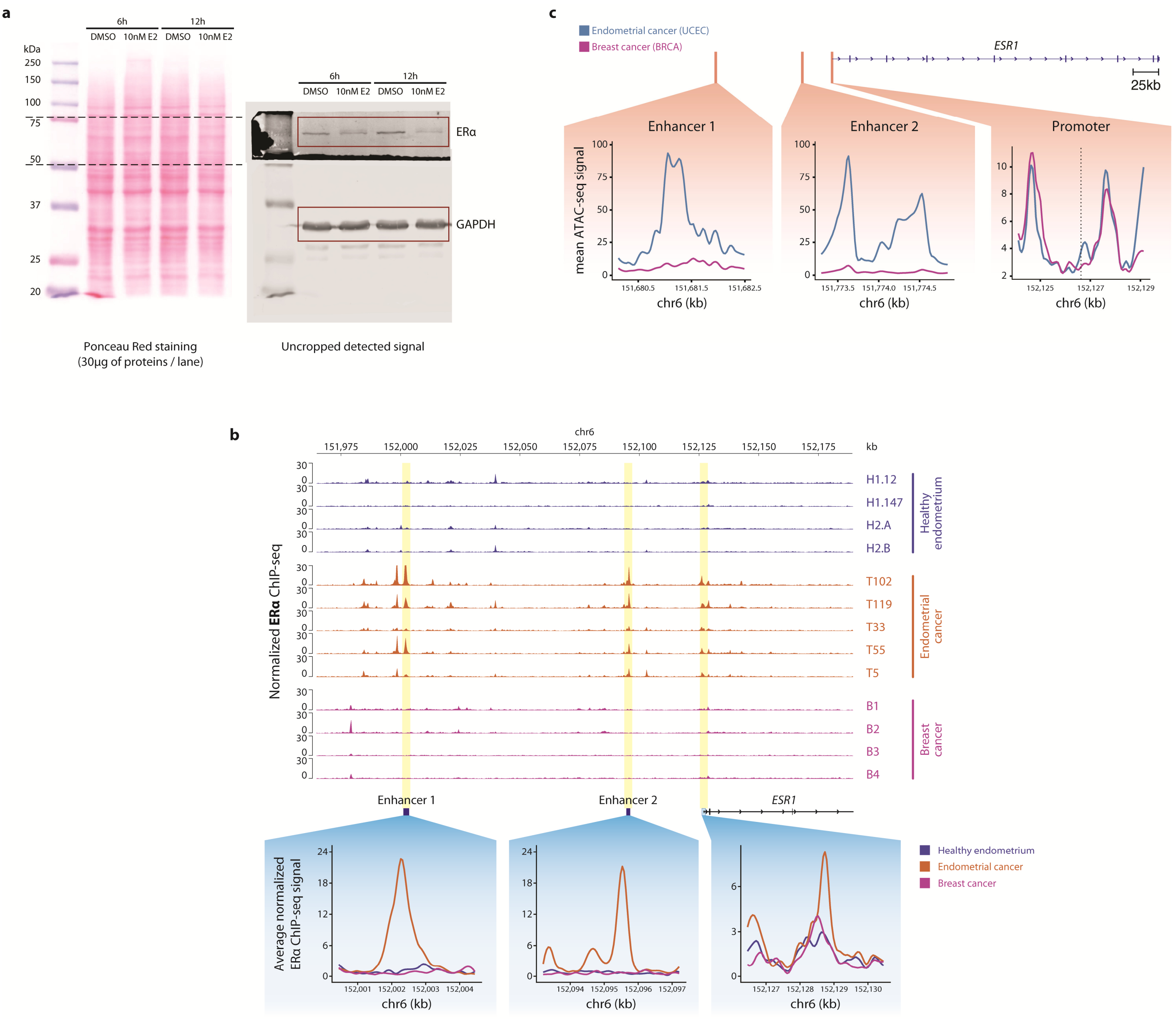


**Supplementary Fig. 7: Genomic and epigenomic landscape comparison of the *ESR1* locus in endometrial and breast cancer.**

**a** Source images and Ponceau Red staining of the western blot shown in **Fig. 3l**. Cropped areas in the main figure are indicated by a red rectangle. **b** Average density profile of ATAC-seq signal from publicly available data (TCGA) for endometrial and breast cancer at the *ESR1* locus (*Enhancer 1*, *Enhancer 2* and promoter). Signal was smoothed using the loess regression method. **c** ChIP-seq genomic tracks for ERα in healthy endometrial tissues (top, blue), endometrial tumors (middle, orange) and breast cancer (bottom, pink) at the *ESR1* locus. Focused average density signal at *ESR1 Enhancer 1*, *Enhancer 2* and promoter are plotted on the lower panel. Average density signal was smoothed using the loess regression method.

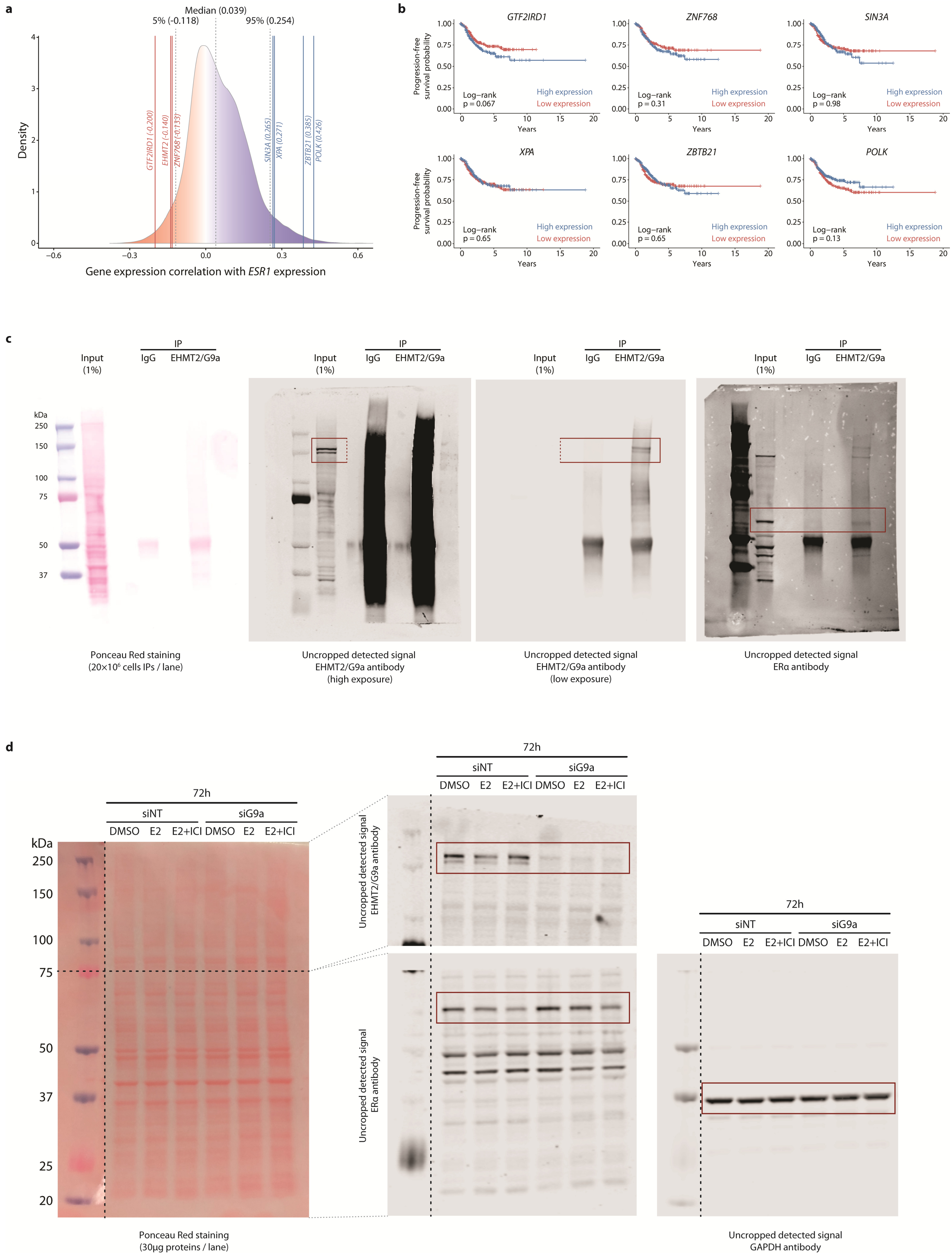


**Supplementary Fig. 8: Validation of the TCGA gene expression analyses.**

**Supplementary Fig. 8: Validation of the TCGA gene expression analyses.**

**a** Density distribution of the *ESR1* gene expression correlation with the whole transcriptome in TCGA RNA-seq data. Vertical blue and red lines indicate the gene expression correlation score between *ESR1* and the top 3 anti-correlated and 3 correlated genes showed in **Fig. 5c**. Black dotted vertical lines indicate respectively the $5^{th}$, $50^{th}$ (median) and $95^{th}$ percentile of the distribution. **b** Progression-Free Kaplan-Meier curve of endometrial cancer patients (TCGA data) divided into two groups using the median of expression gene (FPKM) as cut-off: blue high expression, red low expression. On the upper part are shown the curves for *GTF2IRD1*, *ZNF768*, *SIN3A*, while on the lower part *XPA*, *ZBTB21*, *POLK*. **c-d** Source images and Ponceau Red staining of the western blot shown in **Fig. 5e** (**c**) and **Fig. 5g** (**d**). Cropped areas in the main figure are indicated by red rectangles.

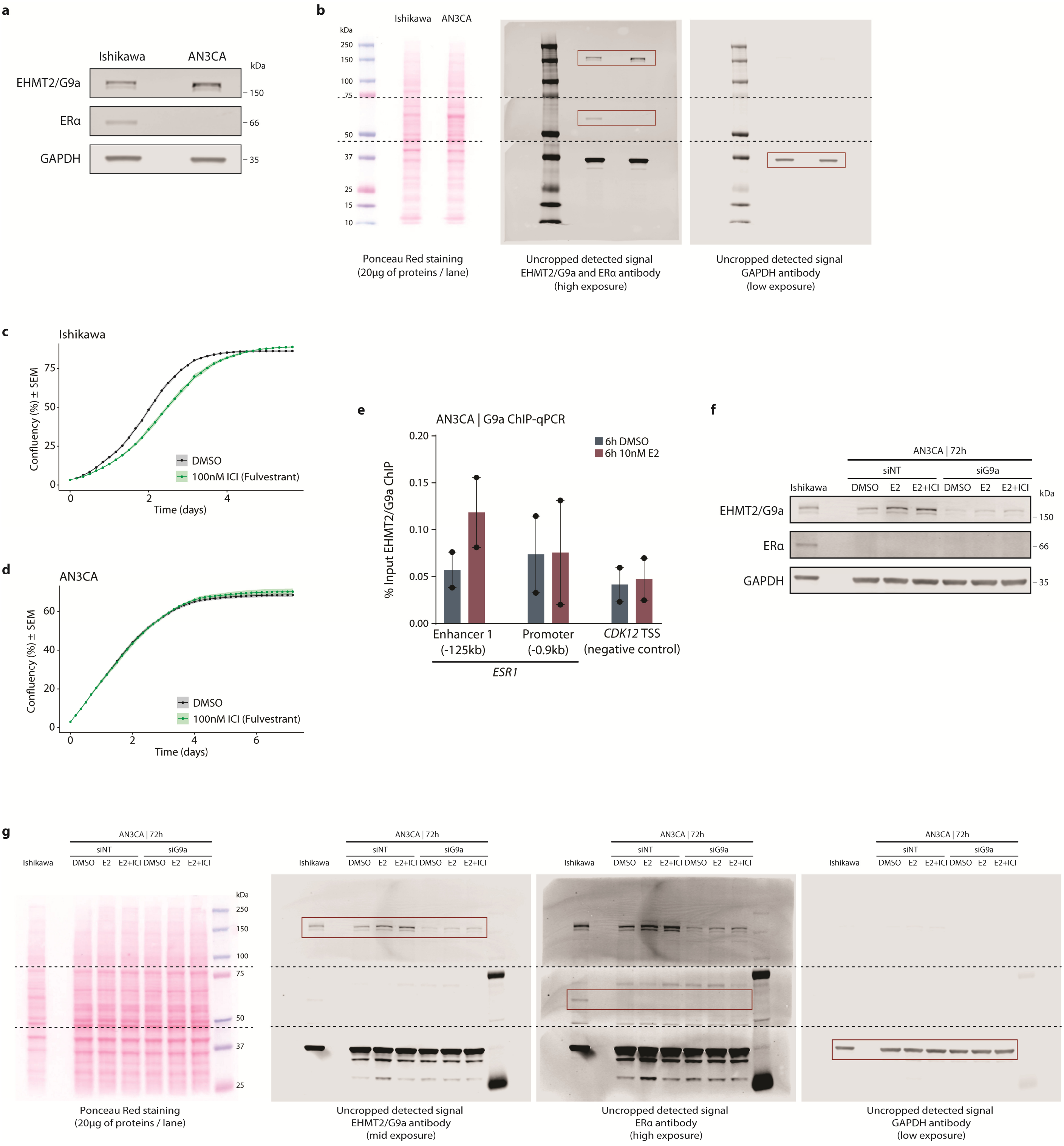


**Supplementary Fig. 9: EHMT2/G9a functional analyses in AN3CA ERα-deficient endometrial cancer cell lines**

**a** Whole protein extracts from Ishikawa and AN3CA endometrial cancer cell lines were immunoblotted using antibodies against EHMT2/G9a, ERα and GAPDH (loading control). **b** Source images and Ponceau Red staining of the western blot shown in (**a**). Cropped areas in the main figure are indicated by red rectangles. **c-d** Normalized cell confluency of Ishikawa (**c**) and *AN3CA* (**d**) endometrial cancer cells upon DMSO (black) or 100nM ICI-182,780/Fulvestrant (green) treatment over time. Average ± SEM of 6 replicates is shown. **e** EHMT2/G9a ChIP in AN3CA cells stimulated for 6h with 10nM β-estradiol. Bar plot shows percentage of enrichment over the input (% Input) at the *ESR1 Enhancer 1*, *ESR1* promoter and *CDK12* promoter (negative control) analyzed by quantitative PCR (qPCR). Mean of 2 independent experiments is shown. **f** AN3CA endometrial cancer cells were stimulated for 72h with 10nM β-estradiol in combination or not with 100nM ICI-182,780 (Fulvestrant). In all conditions, cells where incubated either with a Non-Targeting (NT) siRNA or with an siRNA against *EHMT2/G9a*. Then, whole-cell extracts were analyzed by immunoblotting with antibodies against ERα, EHMT2/G9a and GAPDH (loading control). **g** Source images and Ponceau Red staining of the western blot shown in (**f**). Cropped areas in the main figure are indicated by red rectangles.
